# Abundances of transcripts, proteins, and metabolites in the cell cycle of budding yeast reveals coordinate control of lipid metabolism

**DOI:** 10.1101/2019.12.17.880252

**Authors:** Heidi M. Blank, Ophelia Papoulas, Nairita Maitra, Riddhiman Garge, Brian K. Kennedy, Birgit Schilling, Edward M. Marcotte, Michael Polymenis

## Abstract

Establishing the pattern of abundance of molecules of interest during cell division has been a long-standing goal of cell cycle studies. In several systems, including the budding yeast *Saccharomyces cerevisiae*, cell cycle-dependent changes in the transcriptome are well studied. In contrast, few studies queried the proteome during cell division, and they are often plagued by low agreement with each other and with previous transcriptomic datasets. There is also little information about dynamic changes in the levels of metabolites and lipids in the cell cycle. Here, for the first time in any system, we present experiment-matched datasets of the levels of RNAs, proteins, metabolites, and lipids from un-arrested, growing, and synchronously dividing yeast cells. Overall, transcript and protein levels were correlated, but specific processes that appeared to change at the RNA level (e.g., ribosome biogenesis), did not do so at the protein level, and vice versa. We also found no significant changes in codon usage or the ribosome content during the cell cycle. We describe an unexpected mitotic peak in the abundance of ergosterol and thiamine biosynthesis enzymes. Although the levels of several metabolites changed in the cell cycle, by far the most significant changes were in the lipid repertoire, with phospholipids and triglycerides peaking strongly late in the cell cycle. Our findings provide an integrated view of the abundance of biomolecules in the eukaryotic cell cycle and point to a coordinate mitotic control of lipid metabolism.

## INTRODUCTION

Exemplified by the discovery of cyclin proteins (Evans *et al*., 1983), identifying biomolecules whose abundance changes in the cell cycle has been a critical objective of cell cycle studies for decades. Recognizing such molecular landmarks in the cell cycle is a valuable, and often necessary, step for deciphering how and why cell cycle pathways are integrated.

Over the last twenty years, cell cycle-dependent changes in mRNA levels during the cell cycle of *S. cerevisiae* have been comprehensively defined not only from several arrest-and-release synchronization approaches (Cho *et al*., 1998; Spellman *et al*., 1998; de Lichtenberg *et al*., 2005; Pramila *et al*., 2006; Granovskaia *et al*., 2010), but also elutriation (Spellman *et al*., 1998; Blank *et al*., 2017). Unlike transcript profiling, cell cycle-dependent proteomic and metabolomic changes have been more limited and challenging to interpret due to different or poor synchronization, lack of matched transcriptomic datasets, and divergent results among the various studies. For example, there has only been one mass spectrometry-based proteomic analysis of the budding yeast cell cycle, sampling cultures at four time-points after they were released from arrest (Flory *et al*., 2006). Remarkably few proteins had altered levels during the time course of that experiment, and there was no correlation with the available transcriptomic datasets (Flory *et al*., 2006). Hence, at least in *S. cerevisiae*, it is not clear to what extent protein abundances are dynamic in the cell cycle, and how tightly they are linked to transcriptional changes, if at all.

The picture is not much clearer in other experimental systems. In fission yeast, two recent studies used highly similar arrest-and-release synchronization and protein labeling (stable isotope labeling by amino acids in the cell culture (Mann, 2006)) methods, followed by mass spectrometry, to probe cell cycle-dependent changes in the proteome. In one study only a single protein changed in abundance more than 2-fold (Carpy *et al*., 2014), while in the other report ∼150 proteins did (Swaffer *et al*., 2016). Neither study had experiment-matched transcriptomic datasets. Previously, hundreds of transcripts were reported to be periodic in the cell cycle of fission yeast (Rustici *et al*., 2004; Oliva *et al*., 2005).

In human cells, several reports sampled the proteome in the cell cycle with mass spectrometry, but there is little consensus among them (Dephoure *et al*., 2008; Olsen *et al*., 2010; Lane *et al*., 2013; Ly *et al*., 2014; Becher *et al*., 2018; Dai *et al*., 2018; Schillinger *et al*., 2018). The fraction of proteins identified as periodic ranged from ∼5% (Ly *et al*., 2014), to >65% (Schillinger *et al*., 2018). Synchronization was mostly achieved by release from chemical arrest, but two studies also used elutriation (Ly *et al*., 2014; Dai *et al*., 2018). In the only report where an experiment-matched transcriptomic dataset was generated (Ly *et al*., 2014), the correlation with transcript abundance was positive (ρ=0.63, based on the Spearman rank correlation coefficient). Some of the differences among the above studies may arise from the use of different cell lines, such as: HeLa (Dephoure *et al*., 2008; Olsen *et al*., 2010; Lane *et al*., 2013; Becher *et al*., 2018); K562 (Dai *et al*., 2018); SW480 (Schillinger *et al*., 2018); or NB4 (Ly *et al*., 2014). However, even for the same cell line (HeLa), synchronization (release from thymidine block and nocodazole arrest), and point in the cell cycle (0.5 h after nocodazole arrest), the relative change in abundance of the 3,298 proteins identified in common between the two studies (Olsen *et al*., 2010; Becher *et al*., 2018) was uncorrelated (ρ=0.097, based on Spearman’s rank correlation coefficient; see Materials and Methods).

In *S. cerevisiae*, metabolites have been measured in the cell cycle after arrest-and-release synchronization in minimal medium with ethanol as a carbon source, focusing on exogenous control of cell cycle progression and downstream effects on metabolism (Ewald *et al*., 2016). At the G1/S transition, it is generally thought that cyclin-dependent kinase activity triggers lipolysis (Kurat *et al*., 2009) and mobilizes storage carbohydrates (Ewald *et al*., 2016; Zhao *et al*., 2016), to provide resources for cell division. In other systems, there is evidence of cell cycle-dependent changes on metabolite levels for the green alga *Chlamydomonas reinhardtii* (Juppner *et al*., 2017), fly (Sanchez-Alvarez *et al*., 2015), and human HeLa cells (Atilla-Gokcumen *et al*., 2014; Scaglia *et al*., 2014; Ahn *et al*., 2017). Despite these advances, there has been no experiment-matched sampling of the transcriptome or proteome in any of these studies, making it difficult to integrate these datasets with gene expression, at the mRNA or protein levels.

Here, for the first time in any system, we generated comprehensive datasets for RNAs, proteins, metabolites, and lipids, from the same samples of *S. cerevisiae* cells progressing synchronously in the cell cycle. Importantly, these samples were from elutriated, un-arrested cells, maintaining as much as possible the normal coupling between cell growth and division. We found that while there is a broad correlation between the relative abundances of mRNAs and their corresponding proteins, cell cycle-dependent changes in transcriptional patterns are significantly dampened at the proteome level. The cellular lipid profile is highly cell cycle-regulated, with triglycerides and phospholipids peaking late in the cell cycle, together with protein levels of ergosterol biosynthetic enzymes, highlighting the importance of integrating multiple ‘omic’ datasets to identify cell cycle-dependent cellular processes.

## RESULTS

### Samples for the multi-omic cell cycle analysis

To apply genome-wide methods for the identification of cell cycle-dependent changes in the abundance of molecules of interest, one must first obtain highly synchronous cell cultures. Preferably, synchronization must be achieved in a way that minimally perturbs cellular physiology and the coordination between cell growth and division (Mitchison, 1971; Aramayo and Polymenis, 2017). When cells are chemically or genetically arrested in the cell cycle to induce synchrony, known arrest-related artifacts can bias the results (Mitchison, 1971; Ly *et al*., 2015; Aramayo and Polymenis, 2017). An alternative synchronization method is elutriation, a physical process that fractionates an asynchronous cell population by cell size and sedimentation density properties of the cells, with minimal perturbation of cellular functions (Lindahl, 1948; Creanor and Mitchison, 1979; Banfalvi, 2008). Hence, we used centrifugal elutriation to obtain our synchronous cell cultures (see Materials and Methods, and Figure 1A). Elutriation separates cells primarily based on size, and size is used as a normalizing reference across different elutriation experiments. We isolated 101 different elutriated cultures, which were combined into 24 pools, based on the size at which they were harvested. Hence, we generated a cell size-series, spanning a range from 40 to 75 fL, sampled approximately every 5 fL intervals. These 24 pools were processed as independent samples in all analytical downstream pipelines. For statistical analysis (e.g., with the bootstrap ANOVA), the 24 cell size pools were grouped in 8 groups, for each of the approximately 5 fL increments in the cell size series (see Figure 1A). The same 24 distinct pools were aliquoted as needed (see Materials and Methods) to generate the input samples for measurements of RNA (with RNAseq), proteins (with LC-MS/MS), and metabolites (GC-TOF MS for primary metabolites; HILIC-QTOF MS/MS for biogenic amines; and CSH-QTOF MS/MS for lipids).

**FIGURE 1.**
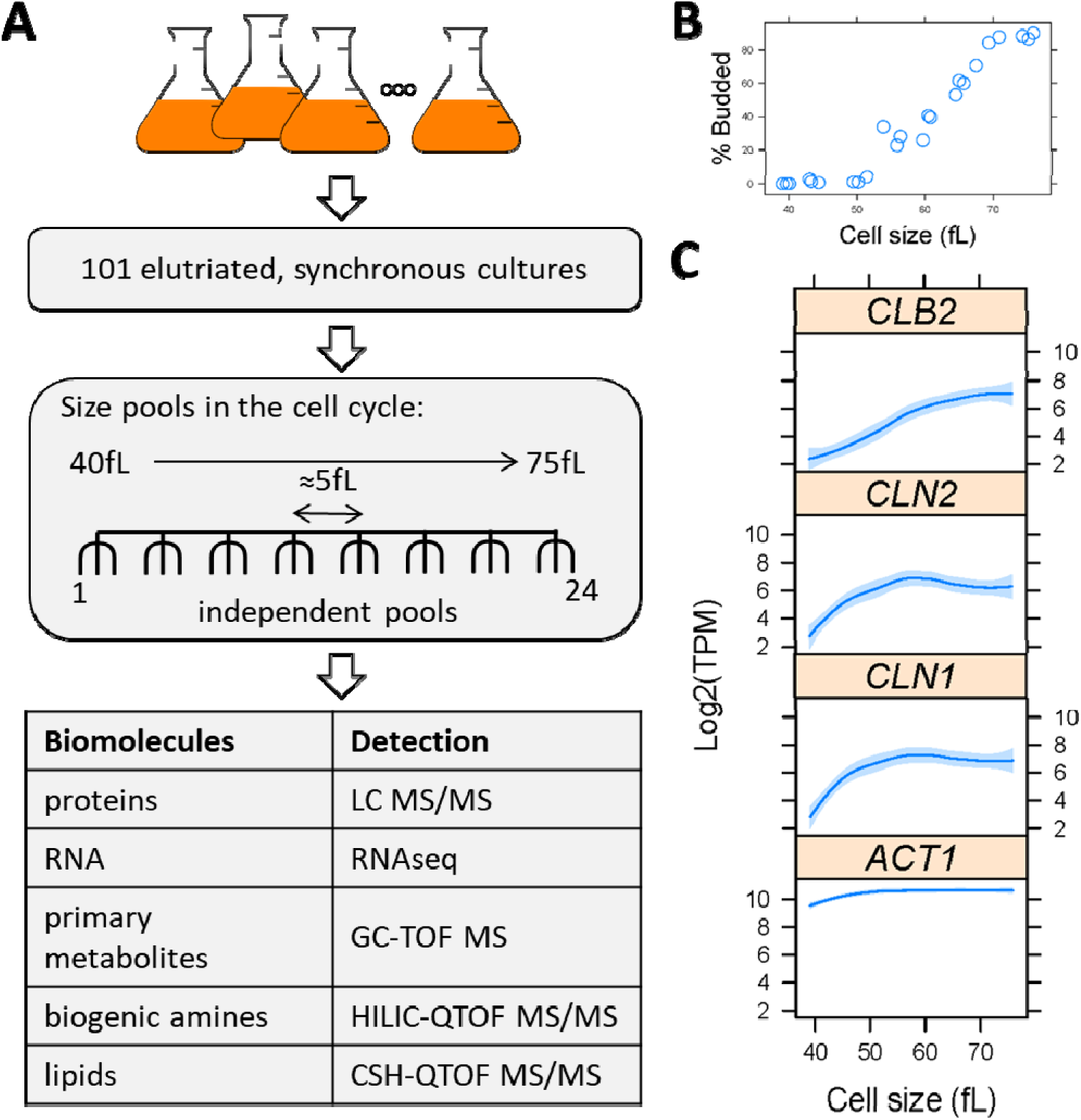
Overview of the experimental design to query cell cycle-dependent changes in the levels of RNAs, proteins, and metabolites. **A**, Generation of sample-matched, multiomic datasets from synchronous cultures of cells of different size, during the cell cycle. **B**, Serving as a morphological marker of cell cycle progression, the percentage of budded cells (y-axis) as a function of cell size (x-axis) is shown for each cell size pool. Cell size corresponds to the mean cell size of the population, and in this case it is the weighted average of all the mean cell sizes of all the elutriated samples that constituted each of the 24 pools. **C**, The levels of mitotic (*CLB2*) or G1 (*CLN1,2*) cyclin mRNAs, which are known to be periodic in the cell cycle, are shown along with those of a non-periodic transcript (*ACT1*; encoding actin). Cell size is shown on the x-axis (in fL), while the Log2-transformed ‘Transcripts Per Kilobase Million’ (TPM) values for each transcript are shown on the y-axis. All 24 values, one for each pool, were plotted in these graphs. Loess curves and confidence bands indicating the standard errors on the curve at a 0.95 level were drawn using the default settings of the panel.smoother function of the *latticeExtra* R language package.

To gauge the synchrony of our samples by microscopy, we used budding as a morphological landmark, which roughly coincides with the initiation of DNA replication in *S. cerevisiae* (Pringle, 1981). The percentage of budded cells across the cell size series (Figure 1B) rose steadily from ∼0% in the smallest cells (at 40 fL), to >80% at the largest cell size (75 fL). The cell size at which half the cells were budded (a.k.a. ‘critical size’, a proxy for the commitment step START) in our cell size series was ∼62 fL (Figure 1B). This value is the same as the critical size these cells display in typical time-series experiments (Hoose *et al*., 2012). We also measured the DNA content of the cells with flow cytometry, confirming the synchrony of the samples (Figure S1). From the RNAseq data that we will describe later (Figure 2), mRNAs that are known to increase in abundance at the G1/S transition (G1 cyclins; *CLN1,2*), or later in G2 phase (cyclin *CLB2*), peaked as expected in the cell size series (Figure 1C). Hence, based on cytological (Figures 1B and S1) and molecular (cyclin mRNAs, Figure 1C) markers of cell cycle progression, the synchrony of our samples was of high quality.

**FIGURE 2.**
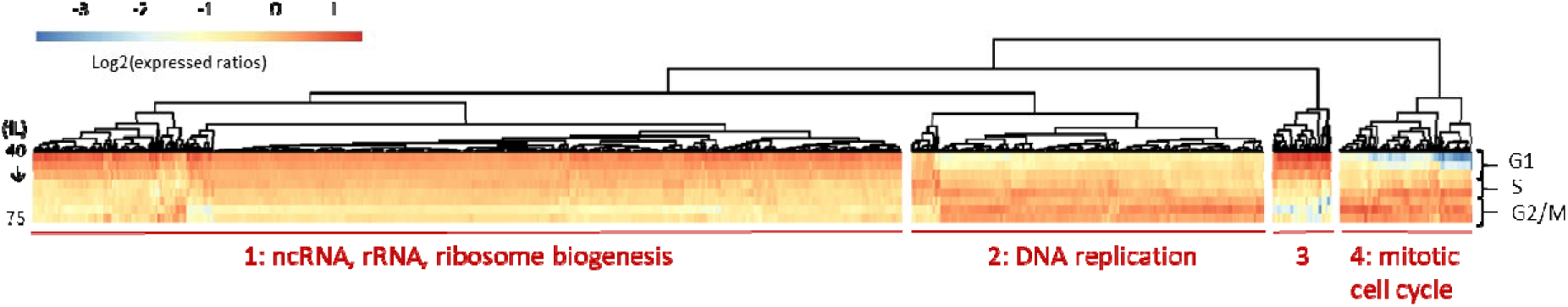
Transcripts changing in abundance in the cell cycle. Heatmap of the levels of 652 differentially expressed RNAs with significantly different levels (p<0.05; Log2(FC)≥1) between any two points in the cell cycle, based on bootstrap ANOVA. The levels of each RNA were the average of each triplicate for the cell size indicated, which was then divided by the average value of the entire cell size series for that RNA. These ‘expressed ratios’ were then Log2-transformed. The Log2(expressed ratios) values were hierarchically clustered and displayed with the *pheatmap* R language package, using the default unsupervised algorithms of the package. The different rows of the heatmap correspond to the different cell sizes (40-75 fL, *top* to *bottom*, in 5fL intervals). The cell cycle phases approximately corresponding to these sizes are shown to the right of the heatmap. The names of all RNAs, values, and clustering classifications are in File4/Sheet: ‘rnas_anova_heatmap’. The gene ontology enrichment analysis for each cluster was done on the PANTHER platform, and the detailed output is in File4/Sheet: ‘rnas_clusters’.

### Overview of the datasets

One type of extract was analyzed for each class of the following biomolecules: RNA, primary metabolites, biogenic amines, and lipids (see Materials and Methods and Table S1). For proteomic analysis, we used soluble protein extracts (designated as ‘sol’ in the datasets, see Table S1) and material from the same extract that was recovered in an insoluble pellet (designated as ‘pel’ in the datasets, see Table S1). The pellet was subsequently solubilized with detergents (see Materials and Methods) and analyzed in parallel to the soluble sample by liquid chromatography tandem mass spectrometry (LC-MS/MS). For label-free relative quantification of proteins, we used both spectral counts (designated as ‘psm’ in the datasets, see Table S1) and peak areas (designated as ‘pa’ in the datasets, see Table S1). For RNAs, the signal we used for quantification was read counts, either raw or after normalization as Transcripts Per kilobase Million (TPM) (see Materials and Methods and Table S1). For the metabolites, the signal was the peak heights from mass spectrometry (designated as ‘ph’ in the datasets, see Table S1). The raw values for all datasets are in File1.

For the quantification of proteins and metabolites, each dataset was first normalized for input. Hence, for proteins or metabolites, comparisons across the 24 samples were scaled based on the sum of the signals detected in each of the 24 samples. For RNA, we used TPM-normalized values and raw reads (see Table S1). All input datasets that entered the downstream computational analyses are in File2. For each dataset, we used a bootstrap-based ANOVA (see Materials and Methods; the output files named as ‘anova’ in the datasets, see Table S1). Also, for RNA, we used the *DESeq2* pipeline ((Love *et al*., 2014); see Materials and Methods; the output file designated as ‘deseq2’, see Table S1). All output datasets are in File3. Only biomolecules that changed ≥2-fold in our cell size series, *and* had an adjusted p-value or FDR<0.05, were considered as significantly changing in the cell cycle.

For display purposes, in all the heatmaps and most plots, we show Log2-transformed expressed ratio values. These are the ratios of the levels that we measured for each biomolecule in each cell cycle point, reflecting the magnitude of the ratio of abundance relative to the average of that biomolecule across all the cell cycle points we sampled. This approach was originally used to describe microarray cell cycle experiments in yeast (Spellman *et al*., 1998), and has been the standard in displaying and analyzing differential expression in the cell cycle.

### RNAs in the cell cycle

The RNAseq data were analyzed (see Materials and Methods, Figure 2, and Table S1), to identify RNAs that change in abundance in the cell cycle. The names of all the RNAs in each set are shown in File4/ Sheet: ‘rna_sets’. The number of identified RNAs varied, depending on the computational method. Based on the *DESeq2* approach, ∼40% of the transcripts (n=2,456) were significantly different between any two points in the cell cycle. The ANOVA-based approach identified 652 RNAs, whose levels changed significantly in the cell size series (Figure 2). In addition to the expected clusters of RNAs associated with DNA replication (cluster 2) and mitotic cell cycle progression (cluster 4), there was a large cluster of transcripts enriched for processes related to ribosome biogenesis (cluster 1, Figure 2; see also File4), peaking in the G1 phase. These transcripts also appeared periodic in past studies that relied on elutriation as a synchronization method to identify cell cycle-regulated RNAs (Spellman *et al*., 1998; Blank *et al*., 2017), but not in studies that used arrest-and-release methods (Spellman *et al*., 1998). An increase in the levels of transcripts involved in ribosome biogenesis before commitment to division has also been described in transcriptomic profiles of *S. pombe* (Oliva *et al*., 2005). Despite these changes at the transcript level, whether the ribosome content of the cell changes during the cell cycle is not known. We will describe results that do not support any cell cycle-dependent changes in assembled ribosomes (Figure 4).

Early in the cell cycle (cluster 1 & 3, Figure 2), we noticed that there were some tRNAs whose levels were higher. Note that tRNAs were not examined in the two prior studies that queried the transcriptome of elutriated *S. cerevisiae* cells, because those studies focused on polyA-tailed selected transcripts (Spellman *et al*., 1998; Blank *et al*., 2017). It has been argued that polyA selection biases the transcriptome quantification (Weinberg *et al*., 2016). Hence, in this study, we relied only on rRNA subtraction to prepare the RNAseq libraries (see Materials and Methods), which does not remove tRNAs and other non-coding RNAs. We also note that tRNAs are notoriously difficult to measure by RNAseq due to factors such as their high level of modification, sequence similarity between different tRNAs, and the difficulty to discriminate between cleaved and mature tRNAs. The tRNAs whose levels appeared to change in the cell cycle are shown in Figure S2. These results are difficult to reconcile with the extreme stability of mature tRNAs (from 9 h to several days-exceeding the duration of multiple cell cycles, see (Hopper, 2013)), unless these tRNAs are targets of quality control mechanisms (Hopper, 2013). In any case, as we show later (Figure S6) we found very little evidence to support a significant role for altered codon usage in the cell cycle.

### Cell cycle-dependent changes in the proteome

From the soluble and insoluble extracts (see Materials and Methods), we identified 3,571 *S. cerevisiae* proteins, at one or more cell cycle points. Although this represents a reasonably thorough sampling of the yeast proteome, we did not find some low abundance proteins (e.g., cyclins). This was not unexpected, since a recent, aggregate analysis of all available datasets of protein abundances in yeast (measured with tandem affinity purification (TAP), followed by immunoblot analysis-, mass spectrometry-, and GFP tag-based methods), placed proteins of the gene ontology process ‘mitotic cell cycle regulation’ as the least abundant group (Ho *et al*., 2018). The extent to which mRNA levels can explain protein levels is debated (Lu *et al*., 2007; Vogel and Marcotte, 2012; Csardi *et al*., 2015; Lahtvee *et al*., 2017). For most species, RNA levels explain between one to two-thirds of the variation in protein abundances (Vogel and Marcotte, 2012). To examine the broad correlation between transcript and protein levels, we looked at the association of count data from our transcriptomic (reads) and proteomic (spectral counts) datasets (Figure S3). Across all the points in our cell size series, the Spearman rank coefficients (ρ) for the transcriptome-proteome correlations ranged from 0.52 to 0.63 (Figure S3).

To identify proteins that changed in abundance in the cell cycle, we examined separately each of the four proteomic datasets: soluble and insoluble extracts, each quantified by spectral counts and by peak areas (see Table S1 and Materials and Methods). The overlap between the proteins in each dataset that appeared to change in abundance in the cell cycle was minimal (see Figure S4). Based on ANOVA analysis, we identified 333 proteins whose levels changed significantly in the cell size series, in at least one of the four proteomic datasets (shown in the heatmap, in Figure 3B). We will describe additional proteins whose levels change significantly in the cell cycle, but due to irregular patterns and missing values were not identified as such by the ANOVA-based method we used (see Figure 5).

**FIGURE 3.**
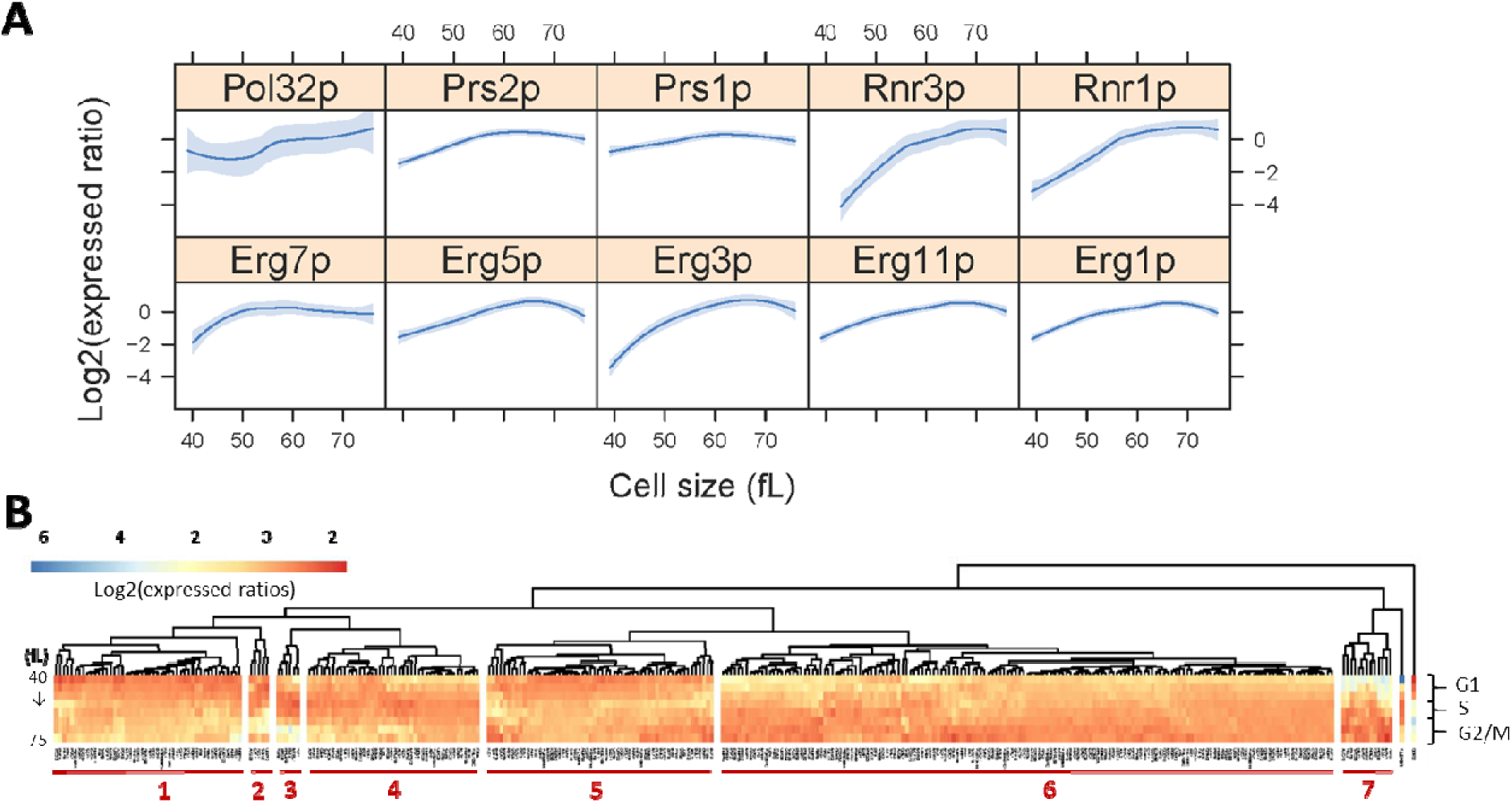
Proteins with cell cycle-dependent abundance. **A**, Levels of selected proteins whose levels changed significantly (p<0.05; Log2(FC)≥1) between any two points in the cell cycle, based on bootstrap ANOVA, in the cell cycle: *Top*, enzymes involved in ergosterol biosynthesis. *Bottom*, enzymes involved in DNA metabolism (Pol32p: DNA polymerase δ; Prs1,2p: PRPP synthase; Rnr1,3p: ribonucleotide-diphosphate reductase). The corresponding Log2(expressed ratios) values from all 24 data points are on the y-axis, and cell size values are on the x-axis. Loess curves and confidence bands indicating the standard errors on the curve at a 0.95 level were drawn using the default settings of the panel.smoother function of the *latticeExtra* R language package. **B**, Heatmap displaying the relative abundance of the 333 proteins in one or more of the four ‘anova’ sets shown in Figure S4. In cases where the same protein was in the intersection of more than one datasets, we chose for display the values from the dataset from which the changes in the protein abundance in the cell cycle was the most significant (i.e., lowest p-value) and greater in magnitude (i.e., highest Log2(FC)). The heatmap was generated as in Figure 2. All the relevant data are in File4/Sheet: ‘proteins_anova_heatmap’.

**FIGURE 4.**
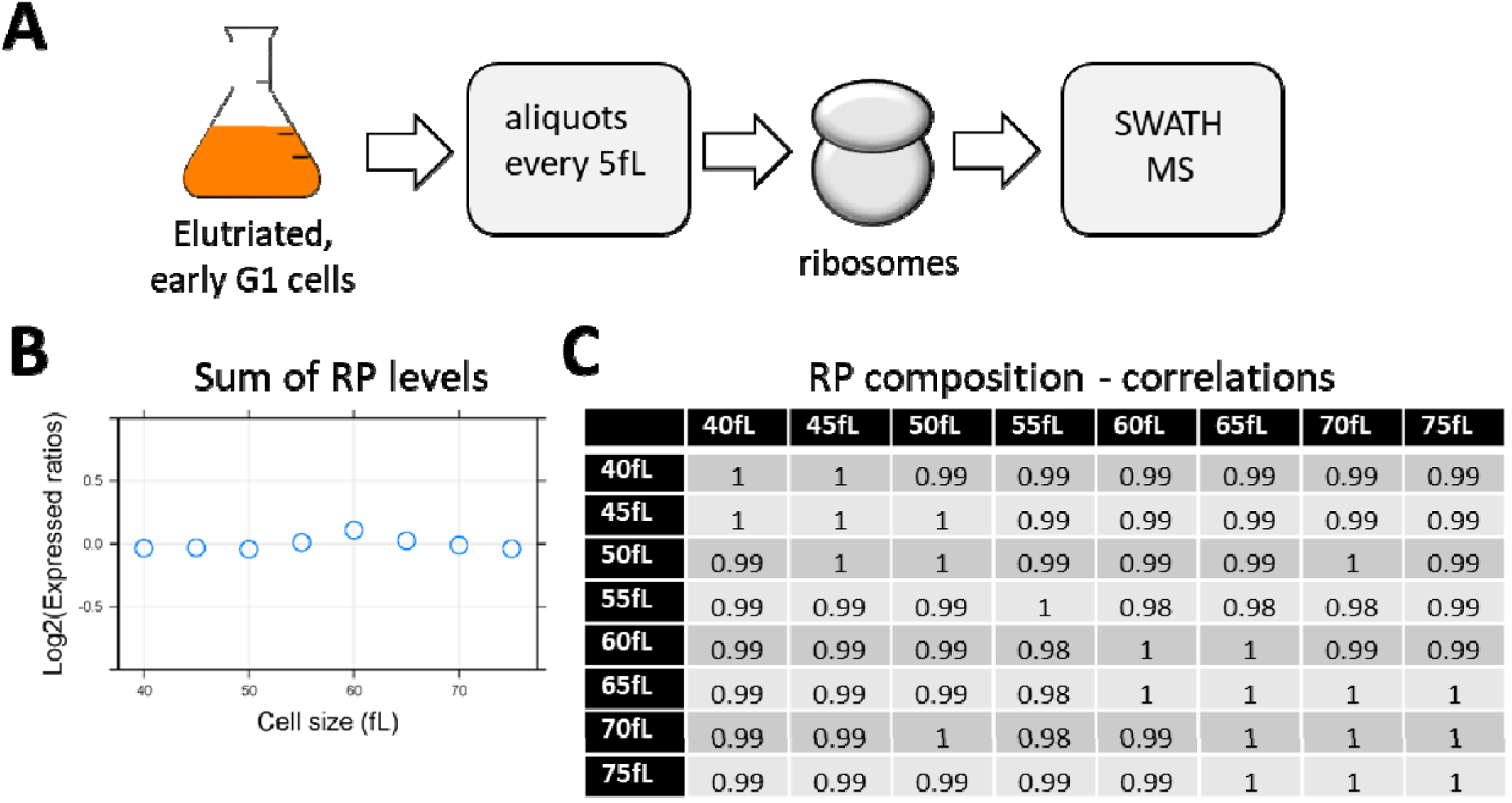
Ribosomal protein abundance in ribosomes does not change in the cell cycle. **A**, Elutriated, early G1 cells were cultured, and sampled at regular intervals in the cell cycle, in three biological replicates at each 5fL range, from 40 to 75 fL. Protein extracts from the same number of cells were then fractionated by sucrose ultra-centrifugation, to isolated ribosomes on mRNAs, which were then analyzed by SWATH-mass spectrometry (see materials and Methods). **B**, The peak areas corresponding to each ribosomal protein (RP) detected were summed and averaged across the triplicate for each cell size interval. The Log2(expressed ratios) values for the ‘Sum of RP levels’ are shown on the y-axis, while cell size is on the x-axis. **C**, Correlation matrix of the relative abundance of individual ribosomal proteins in assembled ribosomes on mRNAs. The Spearman correlation coefficients (ρ) shown in each case were calculated with the rcorr function of the *Hmisc* R language package. The cell cycle profiles for each ribosomal protein are shown in Figure S4.

**FIGURE 5.**
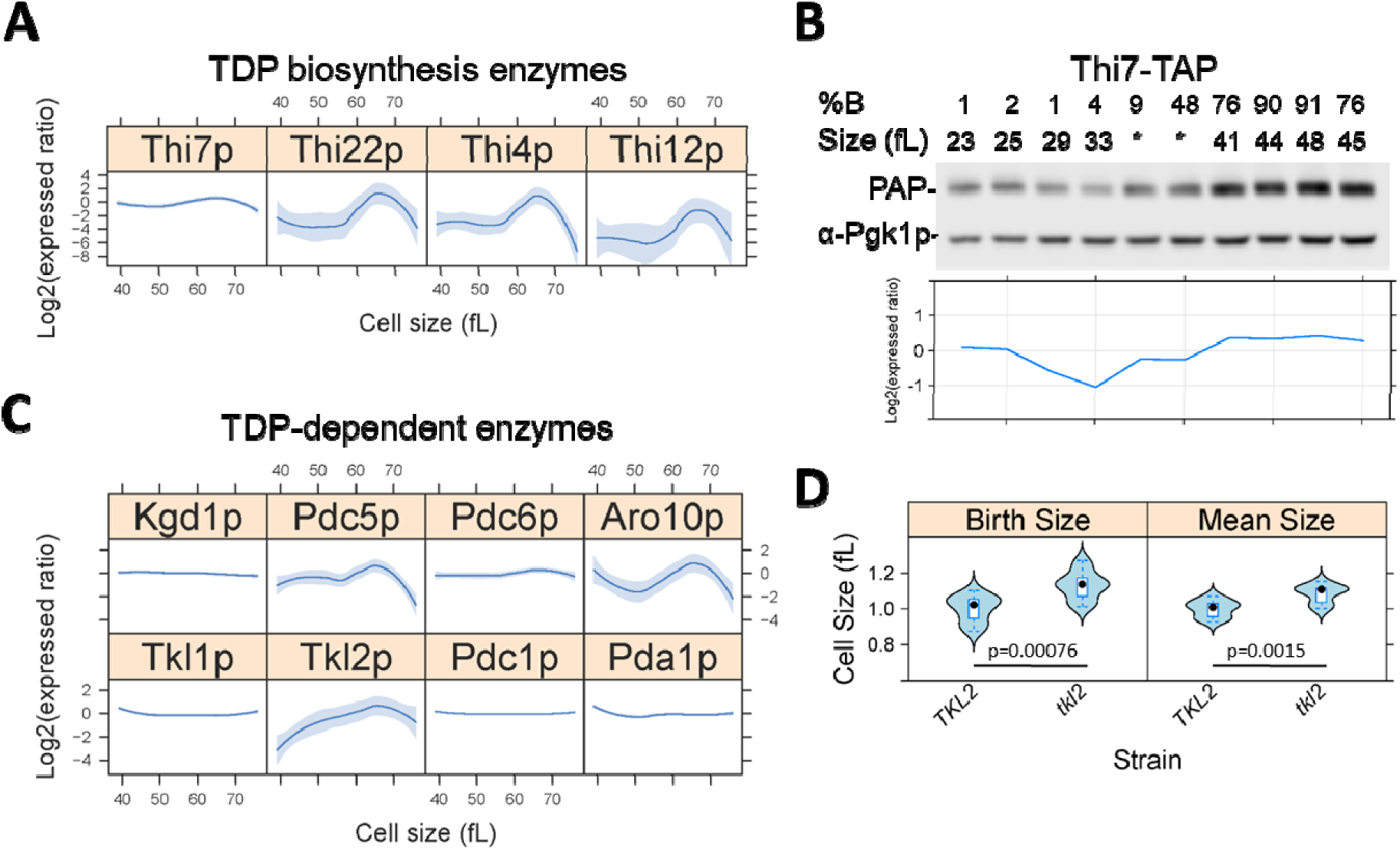
Thiamine biosynthesis and TDP-dependent enzymes in the cell cycle. **A**, Abundances of the indicated proteins of thiamine biosynthesis from LC-MS/MS, across the cell size series (x-axis, in fL). The corresponding Log2(expressed ratios) values from all 24 data points are on the y-axis. Loess curves and confidence bands indicating the standard errors on the curve at a 0.95 level were drawn using the default settings of the panel.smoother function of the *latticeExtra* R language package. **B**, The abundance of Thi7-TAP by immunoblotting from synchronous, elutriated cells, progressing in the cell cycle and sampled at regular intervals, as indicated (%B is the percentage of budded cells; fL is the cell size). Pgk1p levels are also shown from the same samples, to indicate loading. For the two samples indicated with asterisk (*) in the Thi7-TAP series, there were no size data due to instrument malfunction. At the bottom, the band intensities were quantified with ImageJ software, and the Log2-transformed expressed ratios of Thi7-TAP are shown, after they were normalized against Pgk1p. **C**, Abundances of the indicated TDP-dependent proteins, determined and displayed as in A. **D**, The birth and mean size of *tkl2* cells and experiment-matched wild type (*TKL2*) cultures from exponentially dividing cells in rich, undefined media (YPD). At least twelve independent cultures were measured in each case. Significant differences and the associated p values were indicated by the non-parametric Wilcoxon rank sum test, performed with the wilcox.test function of the R *stats* package.

Our analysis provided numerous examples of physiologically relevant, cell cycle-dependent changes in protein abundance. Among these, were several whose levels are well known to be periodic at both the protein and RNA levels. These include proteins involved in DNA replication-related processes, such as both isoforms (Rnr1p and Rnr3p) of the large subunit of ribonucleotide-diphosphate reductase, peaking as cells enter S phase (Figure 3A, bottom). However, other groups of proteins that we found to change in abundance in the cell cycle, were not so at the RNA level. For example, several enzymes of ergosterol biosynthesis (Erg1,11,3,5,7p) peaked late in the cell cycle (Figure 3A, top). Of those, only the levels of the mRNA for Erg3p (C-5 sterol desaturase) changed in the cell cycle (see File4/Sheet: ‘rnas_anova_heatmap’). The coordinate upregulation in the levels of enzymes involved in ergosterol biosynthesis is consistent with the mitotic increase in lipid levels that we will describe later (Figure 6).

**FIGURE 6.**
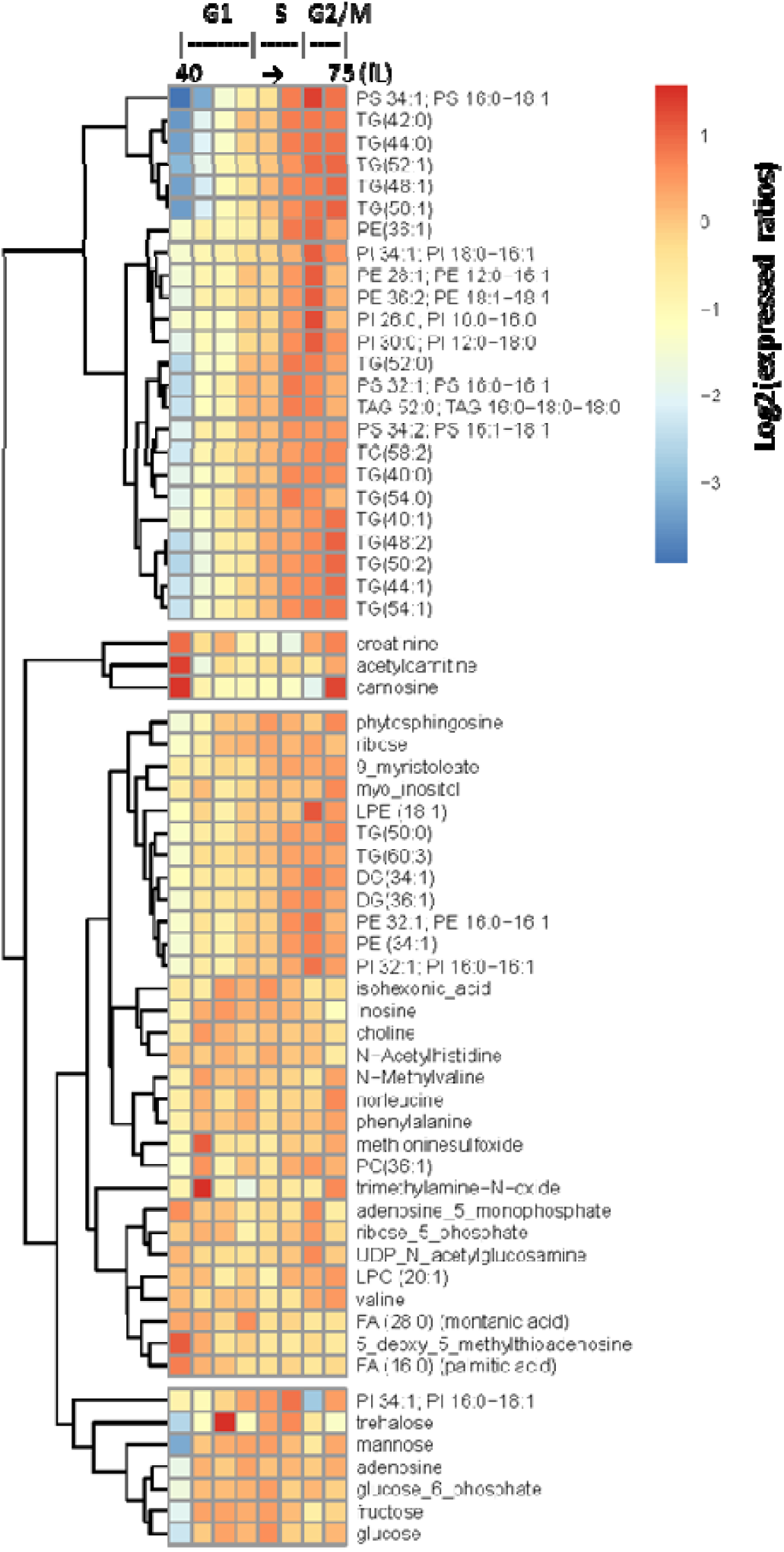
Lipid levels change significantly in the cell cycle. **A**, From 406 known metabolites identified from all classes (primary, biogenic amines, complex lipids), the levels of 64 with significantly different levels (p<0.05; Log2(FC)≥1) between any two points in the cell cycle, based on bootstrap ANOVA, are shown in the heatmap. The levels of each metabolite were the average of each triplicate for the cell size indicated, which was then divided by the average value of the entire cell size series for that metabolite. These ‘expressed ratios’ were then Log2-transformed. The Log2(expressed ratios) values were hierarchically clustered and displayed with the *pheatmap* R language package. The different columns of the heatmap correspond to the different cell sizes (40-75 fL, *left* to *right*, in 5fL intervals).

Despite the transcriptional upregulation in G1 of transcripts involved in ribosome biogenesis (see Figure 2), we did not observe such broad changes at the proteomic level. In earlier reports, the synthesis of ribosomal components was not cell cycle-dependent (Shulman *et al*., 1973; Elliott *et al*., 1979; Warner, 1999). To our knowledge, however, it is not known if the ribosome content in the cell, or the composition of ribosomal proteins in assembled ribosomes, changes in the cell cycle. Hence, we asked if the total amount of ribosomal proteins or their proportion in assembled ribosomes varies significantly in the cell cycle. To this end, we isolated assembled ribosomes through sucrose ultra-centrifugation from wild type cells (Figure 4A; see Materials and Methods). Ribosomal protein abundance was measured with SWATH-mass spectrometry (see Materials and Methods). Note that for this experiment, extracts were not made from pools of different elutriated cultures, but from the same early G1 elutriated cells at different points as they progressed in the cell cycle (see Materials and Methods). Neither the sum of all ribosomal protein abundances (Figure 4B) nor the relative abundance of the individual ribosomal proteins were significantly different in the cell cycle (Figures 4C and S5). These results do not support, but also do not unambiguously exclude, the possibility that individual, specialized ribosomes may be formed during the cell cycle. However, at least based on these population-averaged measurements, ribosome levels and the composition of assembled ribosomes seem unaffected in the cell cycle.

Lastly, we interrogated our proteomic data for evidence of differences in codon usage during the cell cycle. It has been proposed that optimal codon usage is more prevalent in mRNAs expressed in the G1 phase of the cell cycle, contributing to the abundance of proteins that peak in G1 (Frenkel-Morgenstern *et al*., 2012). Altered tRNA abundances during stress conditions in *S. cerevisiae* may also regulate protein synthesis (Torrent *et al*., 2018). To avoid confounding effects from differential transcription of RNAs encoding the proteins that we identified to change in abundance in the cell cycle (Figure 3B), we focused on the proteins whose corresponding mRNAs were not changing in the cell cycle (Figure 2). Moreover, to minimize effects from regulated proteolysis, we excluded from the analysis proteins for which there is evidence for ubiquitylation and regulated proteolysis (Swaney *et al*., 2013). For the vast majority of codons in the remaining proteins, there were no significant changes between their actual and expected frequencies in the cell cycle, based on gene-specific codon usage (Tumu *et al*., 2012). Only four codons (AGC, UAU, AGG, AAC) were used with statistically significant differences in the cell cycle, but the magnitude of those differences was minimal nonetheless (Figure S6). Overall, despite hints at the transcriptional level (Figure 2) for upregulation of processes associated with protein synthesis in the G1 phase, at least from these population-based experiments, our data argue against any significant cell cycle-dependent changes in the ribosome content (Figure 4B), composition (Figure 4C), or codon usage (Figure S6), suggesting that at the proteome level those changes in RNA levels have been dampened extensively.

### Thiamine biosynthesis and TDP-dependent enzymes in the cell cycle

To identify other proteins whose levels could change in the cell cycle but were not identified as such by the computational methods we used, we looked at proteins with the largest change in their levels, regardless of missing values or statistical cutoffs. Remarkably, a group of enzymes involved in thiamine biosynthesis peaked coordinately in abundance late in the cell cycle when the cells reached a cell size of ∼65 fL (Figure 5A). These enzymes participate in thiamine diphosphate (TDP) synthesis in the cytoplasm. To validate these results, we queried in the cell cycle the levels of a TAP-tagged version of Thi7p from a commercially available strain collection (Ghaemmaghami *et al*., 2003), expressed from its endogenous chromosomal location. Thi7p showed the smallest difference (slightly over 2-fold) in abundance during the cell cycle from our mass spectrometry experiments and could provide a good measure to validate our results. Early G1 cells carrying the *THI7-TAP* allele (the only available *THI* gene in the TAP-tagged strain collection encoding any of the proteins shown in Figure 5A) were obtained by elutriation and the levels of the corresponding proteins were evaluated by immunoblotting at regular intervals, as the cultures progressed in the cell cycle (Figure 5B). We confirmed by immunoblotting that the abundance of Thi7p was elevated late in the cell cycle (see Figure 5B; compared to the levels of the control protein Pgk1p). These results are consistent with the notion that there might be a coordinate, mitotic upregulation of thiamine biosynthesis enzymes.

Next, we asked if any TDP-dependent enzymes also change in abundance in the cell cycle and if strains lacking these proteins have cell cycle-related phenotypes. TDP is a cofactor for several enzymes, including transketolase (Tkl1,2p), α-ketoglutarate dehydrogenase (Kgd1p), E1 subunit of pyruvate dehydrogenase (Pda1p), pyruvate decarboxylase (Pdc1,5,6p), and phenylpyruvate decarboxylase (Aro10p). Only the levels of Tkl2p, Pdc5p, and Aro10p appeared to be elevated late in the cell cycle (Figure 5C), at the same time as the levels of thiamine biosynthesis enzymes were also raised (Figure 5A).

Cell size phenotypes are often used as a proxy for disrupted cell cycle progression with an increased cell size phenotype typically accompanying mitotic defects. Of all deletion strains lacking a protein that requires TDP as a cofactor, only the loss of Tkl2p increased cell size significantly (Figure 5D). We found that both birth size and the mean size of *tkl2*Δ cells were larger (Figure 5D). Note that the *tkl2*Δ deletion strain was not in the panels that were examined in genome-wide screens of cell size mutants (Jorgensen *et al*., 2002; Zhang *et al*., 2002). The mitotic upregulation in the levels of thiamine biosynthesis enzymes (Figure 5A) and Tkl2p itself (Figure 5C) are suggestive of possible mitotic roles for Tkl2p, which might depend on the available TDP pools in the cell. In the Discussion, we speculate on such putative roles, based on the published reports.

### Cell cycle-dependent changes in metabolites and lipids

From the same elutriated pools we used to measure RNAs and proteins (see Figure 1), we also measured metabolites and lipids. The assays were performed at the West Coast Metabolomics Center at UC Davis, an NIH RCMRC (Regional Comprehensive Metabolomics Resource Core). Each class of metabolites was measured with distinct mass spectrometry-based assays (see Materials and Methods). From these assays, thousands of compounds were detected, but most could not be assigned confidently to known metabolites, and they were not considered further. Instead, we focused on the 406 primary metabolites, biogenic amines, and complex lipids that were identified across the cell size series. As with our analysis of RNAs and proteins, we used ANOVA (see Table S1 and Figure 6) to identify compounds whose levels change in the cell cycle. Previous reports showed that storage carbohydrates are mobilized at the G1/S transition (Ewald *et al*., 2016; Zhao *et al*., 2016). Consistent with these studies, we also found that trehalose levels rise in G1 to their highest levels when cell size reaches 50 fL, but drop significantly at the G1/S transition (Figure 6). By far, however, the class of metabolites that changed the most in abundance in the cell cycle was complex lipids, which peaked late in the cell cycle (Figure 6). These included phospholipids (phosphatidyl-inositol (PI), -ethanolamine (PE), -serine (PS)) and triglycerides (Figure 6). The higher triglyceride levels are also consistent with the elevated levels of neutral lipid droplets late in the G2/M phase, as reported previously (Blank *et al*., 2017). Overall, the coordinate increase in the levels of ergosterol biosynthesis enzymes we identified from the proteomic analysis (Figure 3A) and the increase in lipids (Figure 6), strongly suggest that lipid metabolism is significantly upregulated late in the cell cycle. In the Discussion, we will expand on the significance of these results.

## DISCUSSION

The sample-matched datasets for RNAs, proteins, metabolites, and lipids we generated from budding yeast cells progressing synchronously in the cell cycle provide a comprehensive view of these biomolecules in dividing cells. We discuss our findings in the context of the relation between the transcriptome and the proteome and the integration of metabolite and lipid measurements with other ‘omic’ datasets.

In yeast, the latest meta-analyses from all available studies estimated that between 37% and 56% of the variance in protein abundance is explained by mRNA abundance (Ho *et al*., 2018). These estimates are within the range of previous ones from multiple species (Vogel and Marcotte, 2012). Based on the absolute quantification of protein and mRNA abundances (Lahtvee *et al*., 2017), the overall correlation between mRNA and protein abundances was also in that range (R^2^=0.45, based on Pearson’s correlation coefficient). The level of correlation between the transcriptome and the proteome we observed appears to be somewhat higher (ρ=0.52-0.63, based on Spearman’s coefficient), probably because our experiments were done from synchronous cells, and because cell cycle transitions are associated with transcriptional waves (Spellman *et al*., 1998). A critical role for transcription in shaping the proteome takes place as cells transition in different environments, and during such transitions changes in protein levels were much more highly correlated with the changes in mRNA levels (R^2^>0.9) (Lahtvee *et al*., 2017). Hence, the relatively high correlation we observed between the transcriptome and the proteome in the cell cycle is not surprising, and it is probably an underestimate, since some extremely unstable cell cycle regulators whose levels rise as a result of transcription (e.g., cyclins, see Figure 1C), were absent from our proteomic datasets because of their low abundance.

Despite the correlation between the transcriptome and the proteome we discussed above, there were clear groups of transcripts and proteins whose abundance was incongruent. Ribosomal biosynthesis, reflected on the levels of individual ribosomal proteins or assembled ribosomes, was not periodic at the proteomic level (Figures 4 and S5), despite a large G1 transcriptional wave of RNAs involved in this process (Figure 2). We noted that a similar phenomenon was recently reported for the integrated stress response, a well-characterized transcriptional response in yeast involving ∼900 transcripts (Gasch *et al*., 2000), which was not seen at all at the protein level (Ho *et al*., 2018). The observation that the ribosome content of the cell is constant in the cell cycle (Figure 4) suggests that changes in translational efficiency of some mRNAs described previously (Blank *et al*., 2017) are likely due to transcript-specific mechanisms, rather than global changes in the steady-state ribosome content (Lodish, 1974).

The mitotic peak in the levels of TDP biosynthesis enzymes was surprising (Figure 5). The physiological significance of such a change in the levels of these enzymes is unclear. Through some uncharacterized roles, the TDP-dependent transketolase activity is necessary for meiotic progression in mouse oocytes (Kim *et al*., 2012). In bacteria, transketolase participates in chromosomal topology, and *E.coli* cells lacking transketolase are UV-sensitive (Hardy and Cozzarelli, 2005). However, we found that yeast *tkl2*Δ cells are not sensitive to UV or other DNA-damaging agents (not shown). Overall, despite the intriguing observations that late in the cell cycle, levels of the TDP-dependent Tkl2p transketolase were higher (Figure 5C) and loss of Tkl2p increased cell size (Figure 5D), the molecular mechanism connecting these observations remains to be determined.

The coordinate upregulation of ergosterol biosynthetic enzymes late in the yeast cell cycle (Figure 3), not evident at the RNA level (Figure 2), was unexpected. To our knowledge, there is no prior report of such a response. It should be noted that the lack of cell cycle-dependent changes at the levels of mRNAs encoding the enzymes of ergosterol biosynthetis was seen not only in our dataset, but also in the other datasets aggregated in the Cyclebase 3.0 database for yeast and other organisms (Santos *et al*., 2015). Of the enzymes we show in Figure 3A, only *ERG3* had a rank score of 624, while all others were not periodic (scores >800) (Santos *et al*., 2015). Note that we also found *ERG3* mRNA levels to significantly change in the cell cycle (see File4/Sheet: ‘rnas_anova_heatmap’).

The mitotic rise in the levels of sterol biosynthetic enzymes is significant in the context of our metabolite measurements, showing that lipid levels (especially phospholipids and triglycerides) increased at the same time (Figure 6). Our observations are consistent with several other reports linking lipid metabolism with cell cycle progression and mitotic entry in yeast (Anastasia *et al*., 2012; McCusker and Kellogg, 2012). Levels of triglycerides increase in wild-type cells synchronized in mitosis (Blank *et al*., 2017), storage of triglycerides in lipid droplets is thought to fuel mitotic exit (Yang *et al*., 2016), and lipid-exchange proteins integrate lipid signaling with cell-cycle progression (Huang *et al*., 2018). Note that there have not been analytical measurements of distinct lipids in the cell cycle in yeast. The data we show here are not only consistent with, but also significantly expand the prior studies mentioned above. It is also important to stress that an increase in lipids late in the cell cycle cannot simply be due to a need for cell surface material. We had shown previously that increased lipogenesis does not increase cell size (Blank *et al*., 2017). Hence, the increase in the abundance of lipids likely reflects changes in the composition of membranes or other more specialized, cell cycle-dependent process, not necessarily a simplistic need for more cell surface building blocks.

One also needs to consider the dramatic changes in cellular morphology. Especially during mitosis, when the cell adopts the characteristic hourglass structure. The lipid content must accommodate dynamic changes in membrane curvature. For example, during cytokinesis, it is thought that lipids that confer negative curvature must be deposited on the outer leaflet of the bilayer (Furse and Shearman, 2018). In yeast and human cells, inhibition of de novo fatty acid biosynthesis arrests cells in mitosis (Hasslacher *et al*., 1993; Schneiter *et al*., 1996; Al-Feel *et al*., 2003; Scaglia *et al*., 2014). In human cells, cholesterol synthesis may affect multiple points in the cell cycle. In an earlier report, inhibition of cholesterol synthesis arrested human cells in mitosis (Suarez *et al*., 2002), while in a later report the cells arrested in G1 (Singh *et al*., 2013). Cholesterol’s role in mitosis appears to be complex, not only affecting the distribution of phospholipids in the plasma membrane but also governing the formation of a vesicular network at the midbody during cytokinesis (Kettle *et al*., 2015). Interestingly, ergosterol may have a cell cycle regulatory role in yeast, distinct from its bulk, structural role in membrane integrity (Dahl *et al*., 1987), but that role remains unclear (Gaber *et al*., 1989). Lastly, our results argue for post-transcriptional mechanisms leading to mitotic upregulation of sterol biosynthesis. As to how the differential abundance of the ergosterol biosynthetic enzymes might come about, we note that all the enzymes we show in Figure 3A, including Erg3p, have been shown to be ubiquitinylated (Peng *et al*., 2003; Swaney *et al*., 2013), raising the possibility of regulated proteolysis.

Overall, our data underscore the value of having metabolite measurements along with other ‘omic’ datasets, to strengthen the efforts of identifying physiologically relevant cellular responses. In future work, employing targeted metabolic profiling and flux analysis in the cell cycle will increase our understanding of how the transcriptome and proteome shape dynamic changes in metabolism and how resources are allocated during cell division.

## ACKNOWLEDGEMENTS

This work was supported by NIH grants R01GM123139 to M.P. and grants from the NIH (R01 HD085901, R01 DK110520, R35 GM122480) and Welch Foundation (F-1515) to E.M.M., with additional mass spectrometry research support from the Army Research Laboratory (Cooperative Agreement # W911NF-17-2-0091). We also acknowledge the support from the NCRR shared instrumentation grant 1S10 OD016281 (Buck Institute) and from NIH grant 1U24DK097154 (UC Davis “West Coast Metabolomics Center”).

## DATA AVAILABILITY

The RNAseq data are deposited at GEO (GSE135476). The LC-MS/MS data are deposited at ProteomeXchange (PXD015273). The SWATH-MS data are deposited at ftp://massive.ucsd.edu/MSV000084302/ with the MassIVE ID MSV000084302; it is also available at ProteomeXchange (PXD015345). All other files related to the data and their analyses are provided as supplements to the manuscript.

## AUTHOR CONTRIBUTIONS

MP and EMM conceptualized the project. HMB, OP, EMM, BKK, BS, and MP designed experiments. MP collected the cells and helped in extract preparation. HMB prepared the RNA samples for RNAseq, performed most of the follow-up experiments for thiamine biosynthesis and TPP-dependent enzymes, and analyzed the relevant data. OP prepared the extracts for LC-MS/MS, ran the mass spectrometry experiments, and analyzed the relevant data. NM examined the cell size of some TPP-dependent enzymes. RG helped with extract preparation for the proteomic samples. MP processed the data, performed most of the analysis, and wrote the first draft of the manuscript. All authors were involved in the editing of the manuscript.

## STRUCTURED METHODS

### REAGENTS AND TOOLS TABLE

Where known, the Research Resource Identifiers (RRIDs) are shown.

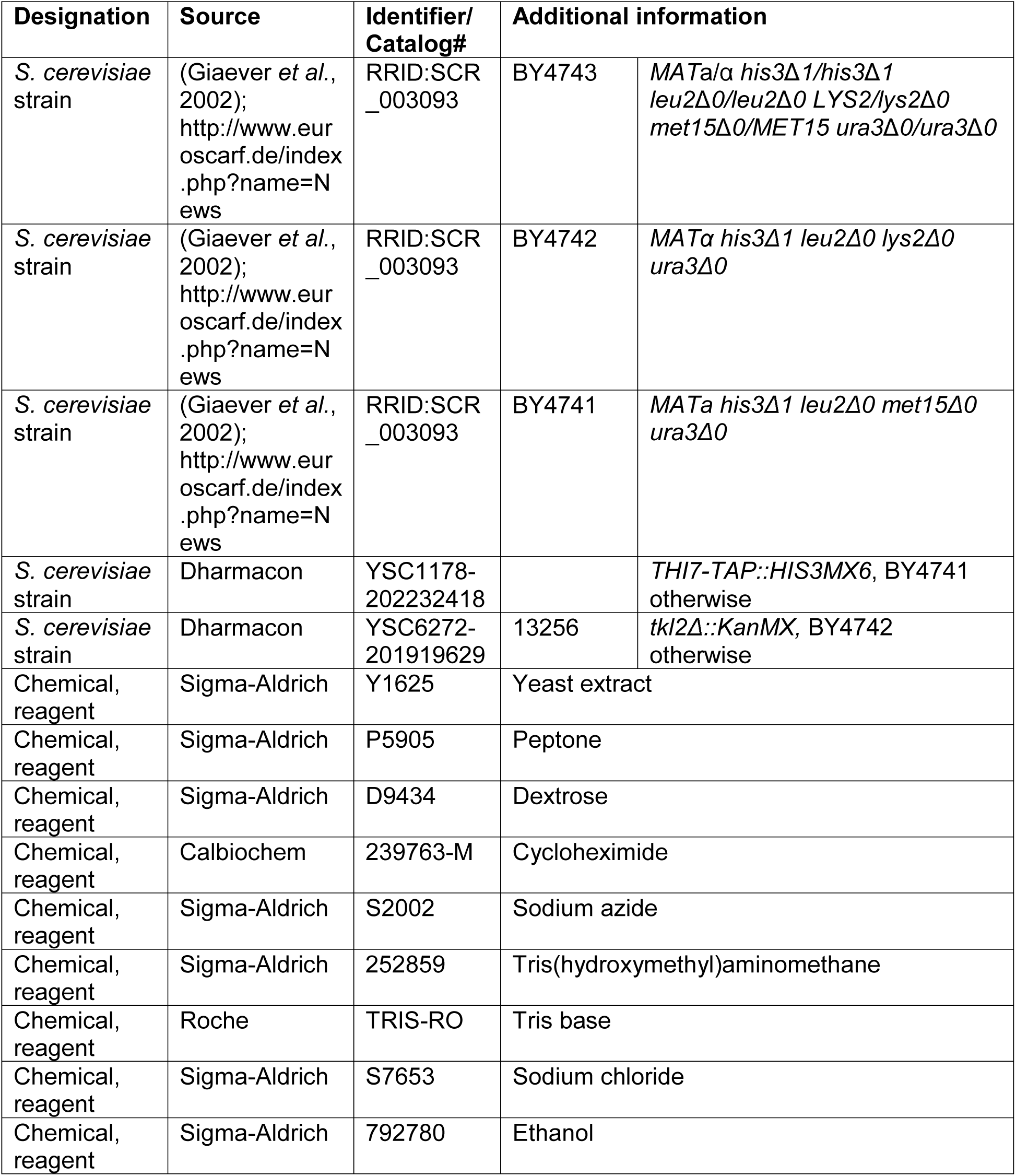
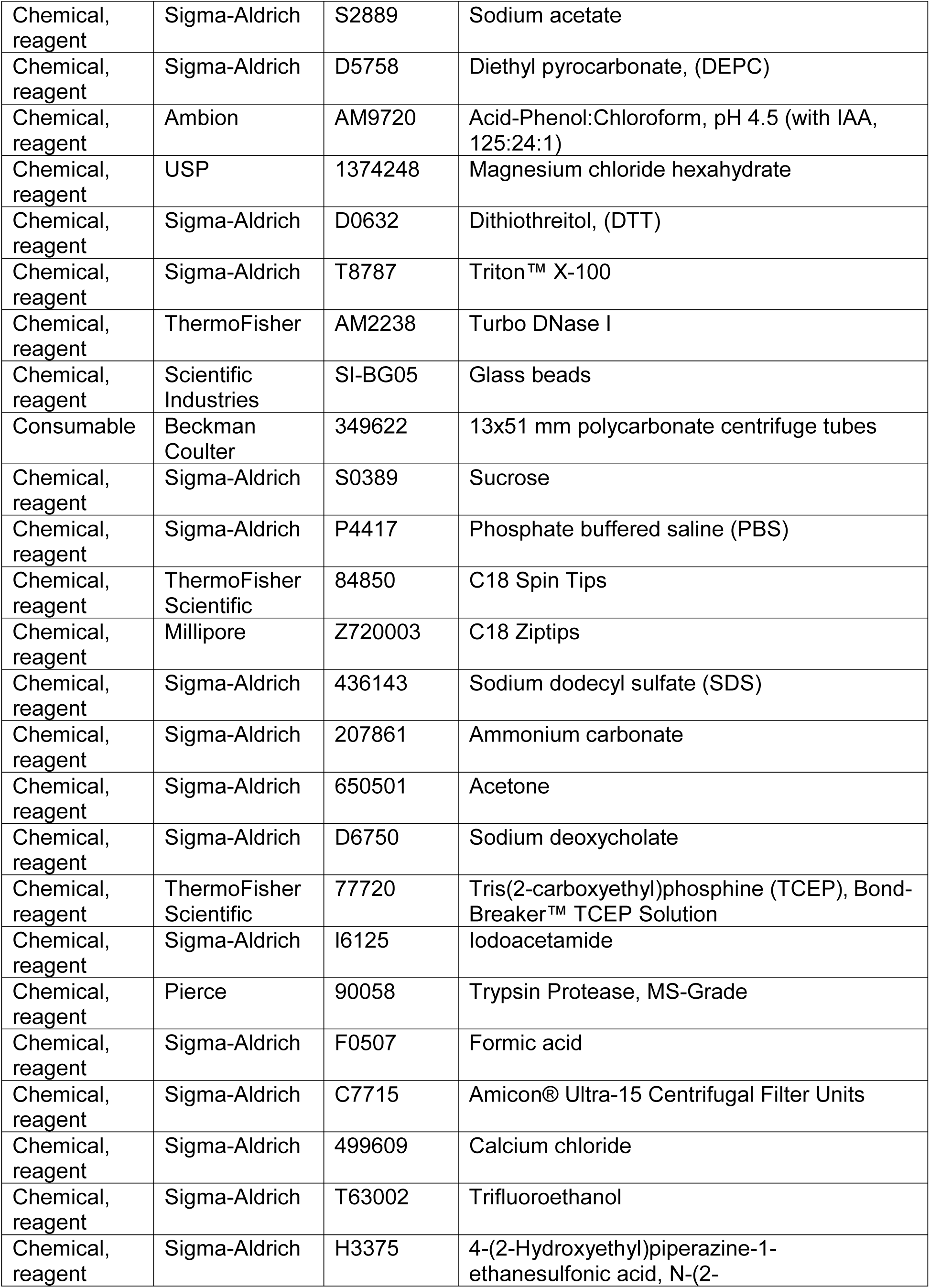
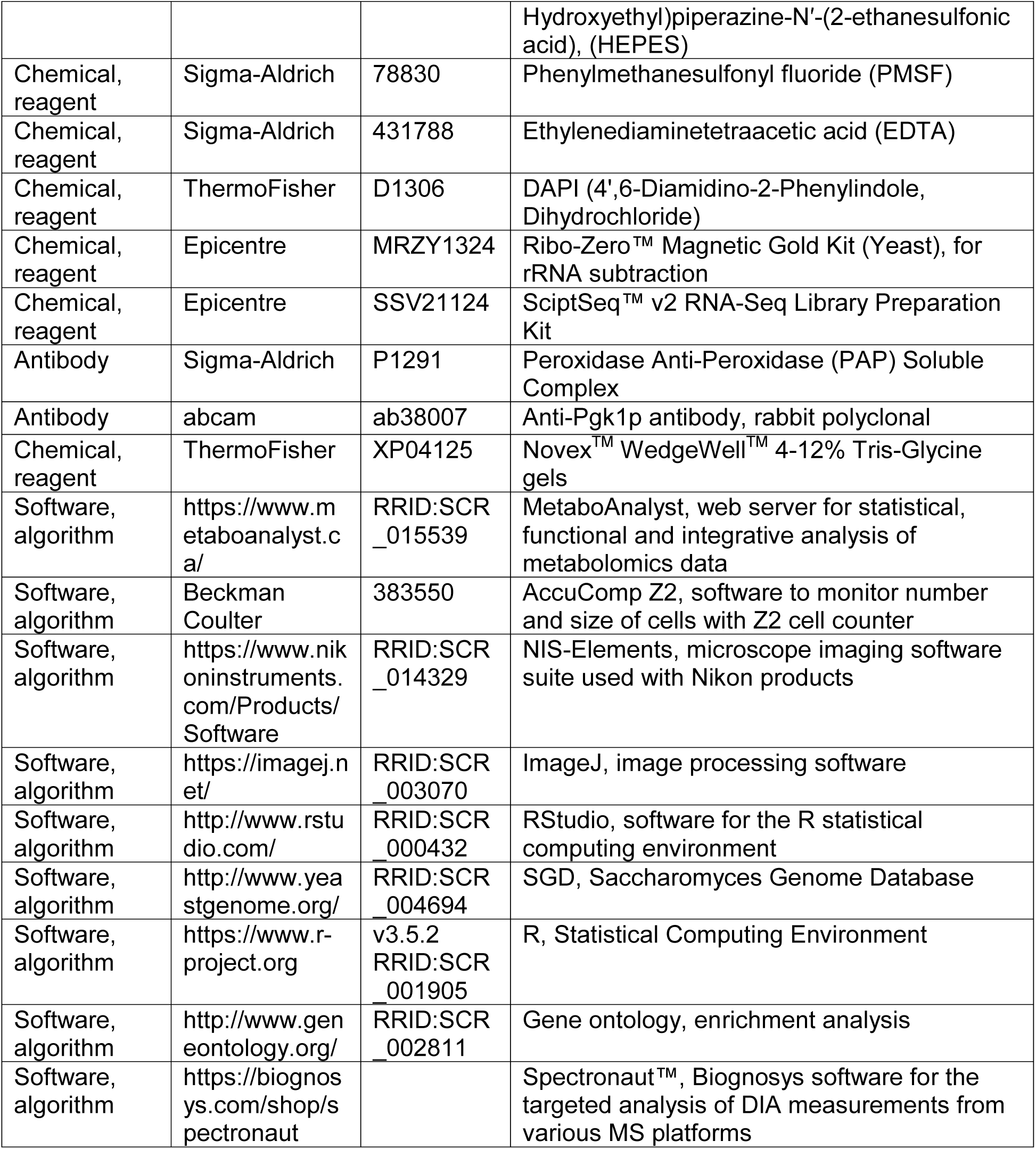

| Designation | Source | Identifier/<br>Catalog# | Additional information |  |
| --- | --- | --- | --- | --- |
| *S. cerevisiae* strain | (Giaever *et al.*, 2002);<br>http://www.euroscarf.de/index.php?name=News | RRID:SCR_003093 | BY4743 | *MATa/α his3Δ1/his3Δ1 leu2Δ0/leu2Δ0 LYS2/lys2Δ0 met15Δ0/MET15 ura3Δ0/ura3Δ0* |
| *S. cerevisiae* strain | (Giaever *et al.*, 2002);<br>http://www.euroscarf.de/index.php?name=News | RRID:SCR_003093 | BY4742 | *MATα his3Δ1 leu2Δ0 lys2Δ0 ura3Δ0* |
| *S. cerevisiae* strain | (Giaever *et al.*, 2002);<br>http://www.euroscarf.de/index.php?name=News | RRID:SCR_003093 | BY4741 | *MATa his3Δ1 leu2Δ0 met15Δ0 ura3Δ0* |
| *S. cerevisiae* strain | Dharmacon | YSC1178-202232418 |  | *THI7-TAP::HIS3MX6*, BY4741 otherwise |
| *S. cerevisiae* strain | Dharmacon | YSC6272-201919629 | 13256 | *tkl2Δ::KanMX*, BY4742 otherwise |
| Chemical, reagent | Sigma-Aldrich | Y1625 | Yeast extract |  |
| Chemical, reagent | Sigma-Aldrich | P5905 | Peptone |  |
| Chemical, reagent | Sigma-Aldrich | D9434 | Dextrose |  |
| Chemical, reagent | Calbiochem | 239763-M | Cycloheximide |  |
| Chemical, reagent | Sigma-Aldrich | S2002 | Sodium azide |  |
| Chemical, reagent | Sigma-Aldrich | 252859 | Tris(hydroxymethyl)aminomethane |  |
| Chemical, reagent | Roche | TRIS-RO | Tris base |  |
| Chemical, reagent | Sigma-Aldrich | S7653 | Sodium chloride |  |
| Chemical, reagent | Sigma-Aldrich | 792780 | Ethanol |  |

|  |  |  |  |
| --- | --- | --- | --- |
| Chemical, reagent | Sigma-Aldrich | S2889 | Sodium acetate |
| Chemical, reagent | Sigma-Aldrich | D5758 | Diethyl pyrocarbonate, (DEPC) |
| Chemical, reagent | Ambion | AM9720 | Acid-Phenol:Chloroform, pH 4.5 (with IAA, 125:24:1) |
| Chemical, reagent | USP | 1374248 | Magnesium chloride hexahydrate |
| Chemical, reagent | Sigma-Aldrich | D0632 | Dithiothreitol, (DTT) |
| Chemical, reagent | Sigma-Aldrich | T8787 | Triton™ X-100 |
| Chemical, reagent | ThermoFisher | AM2238 | Turbo DNase I |
| Chemical, reagent | Scientific Industries | SI-BG05 | Glass beads |
| Consumable | Beckman Coulter | 349622 | 13x51 mm polycarbonate centrifuge tubes |
| Chemical, reagent | Sigma-Aldrich | S0389 | Sucrose |
| Chemical, reagent | Sigma-Aldrich | P4417 | Phosphate buffered saline (PBS) |
| Chemical, reagent | ThermoFisher Scientific | 84850 | C18 Spin Tips |
| Chemical, reagent | Millipore | Z720003 | C18 Ziptips |
| Chemical, reagent | Sigma-Aldrich | 436143 | Sodium dodecyl sulfate (SDS) |
| Chemical, reagent | Sigma-Aldrich | 207861 | Ammonium carbonate |
| Chemical, reagent | Sigma-Aldrich | 650501 | Acetone |
| Chemical, reagent | Sigma-Aldrich | D6750 | Sodium deoxycholate |
| Chemical, reagent | ThermoFisher Scientific | 77720 | Tris(2-carboxyethyl)phosphine (TCEP), Bond-Breaker™ TCEP Solution |
| Chemical, reagent | Sigma-Aldrich | I6125 | Iodoacetamide |
| Chemical, reagent | Pierce | 90058 | Trypsin Protease, MS-Grade |
| Chemical, reagent | Sigma-Aldrich | F0507 | Formic acid |
| Chemical, reagent | Sigma-Aldrich | C7715 | Amicon® Ultra-15 Centrifugal Filter Units |
| Chemical, reagent | Sigma-Aldrich | 499609 | Calcium chloride |
| Chemical, reagent | Sigma-Aldrich | T63002 | Trifluoroethanol |
| Chemical, reagent | Sigma-Aldrich | H3375 | 4-(2-Hydroxyethyl)piperazine-1-ethanesulfonic acid, N-(2- |
|  |  |  | Hydroxyethyl)piperazine-N'-(2-ethanesulfonic acid), (HEPES) |
| Chemical, reagent | Sigma-Aldrich | 78830 | Phenylmethanesulfonyl fluoride (PMSF) |
| Chemical, reagent | Sigma-Aldrich | 431788 | Ethylenediaminetetraacetic acid (EDTA) |
| Chemical, reagent | ThermoFisher | D1306 | DAPI (4',6-Diamidino-2-Phenylindole, Dihydrochloride) |
| Chemical, reagent | Epicentre | MRZY1324 | Ribo-Zero™ Magnetic Gold Kit (Yeast), for rRNA subtraction |
| Chemical, reagent | Epicentre | SSV21124 | ScriptSeq™ v2 RNA-Seq Library Preparation Kit |
| Antibody | Sigma-Aldrich | P1291 | Peroxidase Anti-Peroxidase (PAP) Soluble Complex |
| Antibody | abcam | ab38007 | Anti-Pgk1p antibody, rabbit polyclonal |
| Chemical, reagent | ThermoFisher | XP04125 | Novex™ WedgeWell™ 4-12% Tris-Glycine gels |
| Software, algorithm | https://www.metaboanalyst.ca/ | RRID:SCR_015539 | MetaboAnalyst, web server for statistical, functional and integrative analysis of metabolomics data |
| Software, algorithm | Beckman Coulter | 383550 | AccuComp Z2, software to monitor number and size of cells with Z2 cell counter |
| Software, algorithm | https://www.nikoninstruments.com/Products/Software | RRID:SCR_014329 | NIS-Elements, microscope imaging software suite used with Nikon products |
| Software, algorithm | https://imagej.net/ | RRID:SCR_003070 | ImageJ, image processing software |
| Software, algorithm | http://www.rstudio.com/ | RRID:SCR_000432 | RStudio, software for the R statistical computing environment |
| Software, algorithm | http://www.yeastgenome.org/ | RRID:SCR_004694 | SGD, Saccharomyces Genome Database |
| Software, algorithm | https://www.r-project.org | v3.5.2<br>RRID:SCR_001905 | R, Statistical Computing Environment |
| Software, algorithm | http://www.geneontology.org/ | RRID:SCR_002811 | Gene ontology, enrichment analysis |
| Software, algorithm | https://biognosys.com/shop/spectronaut |  | Spectronaut™, Biognosys software for the targeted analysis of DIA measurements from various MS platforms |

### METHODS AND PROTOCOLS

#### Strains and media

All the strains used in this study are shown in the Key Resources Table, above. Unless noted otherwise, the cells were cultivated in the standard, rich, undefined medium YPD (1% ^w^/_v_ yeast extract, 2% ^w^/_v_ peptone, 2% ^w^/_v_ dextrose), at 30 °C (Kaiser *et al*., 1994).

#### Elutriation

To collect enough cells for the downstream measurements of RNA, proteins, and metabolites, we followed the same strategy we described previously (Blank *et al*., 2017). Briefly, elutriated wild type, G1 cells (diploid BY4743 background) were allowed to progress in the cell cycle until they reached the desired cell size. At that point, they were quenched (with 100 µg/ml cycloheximide and 0.1% sodium azide) and frozen away, and later pooled with cells of similar size (Figure 1A). Overall, we had to collect 101 individual samples, to generate the 24 pools shown in Figure 1A.

For other elutriation experiments (e.g., see Figures 4,5), only an early G1 elutriated fraction was collected, from which samples were taken at regular intervals as the cells progressed in the cell cycle.

#### Cell size and DNA content measurements

The methods to measure DNA content and the cell size (birth or mean size) of asynchronous cultures and estimate the critical size of asynchronous cultures, have been described in detail previously (Guo *et al*., 2004; Truong *et al*., 2013; Soma *et al*., 2014; Maitra *et al*., 2019).

#### Proteomic samples

We used ∼1E+09 cells from each of the 24 pools of the cell size series (see Figure 1) to prepare extracts for LC-MS/MS. For each sample, the cells were resuspended in 0.75 ml of lysis solution (10 mM Tris pH 7.8, 10 mM NaCl). Glass beads were added to the top of liquid level, the samples were placed in a Mini Beadbeater (Biospec), and the cells broken by ‘bead-beating’ twice at the maximum speed for 90 s each time, placed on ice for 60 s between. The extract from each sample was collected by punching a hole with a 21-gauge syringe needle at the bottom of the tube. Lastly, the soluble material from the lysates were clarified by centrifugation at 14,000 *g* at 4 °C, for 10 m. Insoluble pellets were resuspended in 500 μl of lysis buffer and both clarified supernatants and pellets were stored at −80 °C until processing for mass spectrometry.

For mass spectral analysis, clarified extracts were thawed and protease inhibitors immediately added. 50 μl of each supernatant sample was mixed with 50 µl trifluoroethanol (TFE) and reduced with 5mM tris(2-carboxyethyl)phosphine (TCEP) at 56 °C for 45 m, cooled for 5 m at room temperature, and alkylated with 25 mM iodoacetamide in the dark, at room temperature for 30 m. Samples were diluted 10-fold with digestion buffer (50 mM Tris pH 8.0, 2 mM calcium chloride), digested with trypsin (added at 1:50 ratio) at 37 °C for 5 h. Digestion was stopped with 100 µl of 10% formic acid and sample volumes were reduced to 100-250 μl in a SpeedVac. Following filtration with an Amicon® Ultra-15 Centrifugal Filter Unit the peptides were desalted using C18 Spin Tips, according to the manufacturer’s instructions. The volume of the samples was then reduced to 5-10 μl in a SpeedVac. Lastly, the samples were resuspended in 100 μl of a 95% water, 5% acetonitrile, 0.1% formic acid solution, and subjected to LC-MS/MS analysis.

The insoluble pellets from the same extracts described above were processed based on a method reported previously (Lin *et al*., 2013). The pellets were resuspended in 50 μl of 2% ^w^/_v_ sodium dodecyl sulfate (SDS), 50 mM ammonium carbonate and heated at 95 °C for 10 m.

Following clarification each supernatant was transferred to a fresh tube, mixed with six volumes of cold acetone (−20 °C), and incubated at 4 °C for 4 h to form a precipitate. Precipitate was recovered by centrifugation at 13,000 g for 15 m, the supernatant carefully removed by aspiration, and the pellets washed twice with 0.4 ml of cold acetone. After each wash the samples were centrifuged at 14,000 g for 1 m and the supernatant carefully aspirated. Pellets were solubilized in 500 μl of 1% w/v sodium deoxycholate, 50 mM ammonium carbonate with two rounds of sonication (10 m each) in a water bath sonicator with 5 m on ice in between. 50 µl of each sample was reduced and alkylated with TCEP and iodocateamide as described above. Unreacted iodoacetamide was quenched with 12 mM dithiothreitol (DTT). The samples were brought to 80 µl with digestion buffer and digested with trypsin (added at 1:50 ratio) at 37 °C for 5 h. Digestion was stopped with 1% formic acid and samples were centrifuged at 14,000 g for 10 m to pellet the precipitated sodium deoxycholate. Peptides were desalted with C18 Spin Tips, and resuspended for LC-MS/MS as described above.

#### LC-MS/MS

Mass spectra were acquired on a Thermo Orbitrap Fusion. 5 µl (supernatant samples) or 2 µl (pellet samples) of peptides were separated using reverse phase chromatography on a Dionex Ultimate 3000 RSLCnano UHPLC system (Thermo Scientific) with a C18 trap to Acclaim C18 PepMap RSLC column (Dionex; Thermo Scientific) configuration. Peptides were eluted using a 3-45% acetonitrile gradient over 70 min and directly injected into the mass spectrometer using nano-electrospray. Data-dependent tandem mass spectrometry was performed using a top speed HCD method with full precursor ion scans (MS1) collected at 120,000 m/z resolution and a cycle time of 3 sec. Monoisotopic precursor selection and charge-state screening were enabled, with ions of charge >□+□1 selected with dynamic exclusion of 30□s for ions selected once within a 30□s window. Selected precursor ions underwent high-energy collision-induced dissociation (HCD) at 31% energy stepped +/−4%. All MS2 scans were centroid and done in rapid mode. Raw files were processed using Proteome Discoverer 2.2 and the label-free quantification workflow.

#### RNA samples and libraries

We used the same approach we had described previously (Blank *et al*., 2017), to collect cells from elutriated cultures of wild type (BY4743 strain background). For each of the 24 samples, from ∼3E+07 cells total RNA was prepared with the hot phenol method. Briefly, the frozen pellets were re-suspended in 0.4 ml TES buffer (10 mM Tris pH = 7.5, 10mM EDTA, 0.5% SDS), in DEPC-treated water, and ∼0.05 ml glass beads were added. Then, 0.4 ml of acid phenol:chloroform was added to each pellet, and the samples were incubated at 65 °C for 30 m, and vortexed briefly every 5 m during that time. The samples were centrifuged at 14,000 *g* for 5 m, and 0.3 ml of the top, aqueous layer were placed in a 2-ml screw-cap tube containing 1 ml cold ethanol with 40 μl of a 3M sodium acetate solution. The samples were incubated at 4 °C overnight and then centrifuged at 14,000 *g* for 20 m. The pellets were washed with 80% ethanol and centrifuged at 14,000 *g* for 5 m. The pellets were air-dried and resuspended in 25 μl of DEPC-treated water. For the RNAseq libraries, we also used the same approach we had described (Blank *et al*., 2017), except that we did not select for polyA-tailed RNAs. Instead, from total RNA, we depleted rRNA, using the ‘Ribo-Zero™ Magnetic Gold Kit (Yeast)’, according to the manufacturer’s instructions. All libraries were sequenced on an Illumina HiSeq4000, with multiplexing, at the Texas A&M AgriLife Genomics and Bioinformatics Facility. Raw sequencing data (fastq files) have been deposited (GEO: GSE135476).

The reads were aligned to the *S. cerevisiae* reference genome (version R64-1-1) using the *Rsubread* R language package (Liao *et al*., 2019). First, an index was built using the command: buildindex(basename = “R64”, reference = “Saccharomyces_cerevisiae.R64-1-1.dna.toplevel.fa”, gappedIndex=TRUE). Then, for each of the 24 libraries, the paired end reads were aligned with the command: align(index = ‘R64’, readfile1 = ‘….fastq.gz’, readfile2 = ‘….fastq.gz’, type = “rna”). For each library, we obtained >10 million uniquely mapped reads, and the output BAM files were then used in the featureCounts function of the *Rsubread* package, with the following command: featureCounts(files = “…subread.BAM”, ispairedEnd = TRUE, requireBothEndsMapped = TRUE, annotext = “Saccharomyces_cerevisiae.R64-1-1.95.gtf”, countChimericFragments = FALSE, isGTFAnnotationFile = TRUE). All the read counts are in File1/sheet ‘rna_reads’.

For differential RNA levels between any two points in the cell cycle using the *DESeq2* R language package (Love *et al*., 2014), the raw read data (File2/sheet ‘rna_deseq2_i’) were used as input. For this statistical analysis, the 24 cell size pools were grouped in 8 groups, for each of the approximately 5 fL increments in the cell size series (see Figure 1A). Additional analyses with ANOVA-based methods were performed as for the other biomolecules, and they are described below.

#### Metabolite samples and analysis

The untargeted, primary metabolite, biogenic amine, and complex lipid analyses were done at the NIH-funded West Coast Metabolomics Center at the University of California at Davis, according to their mass spectrometry protocols. Gas Chromatography–Time-of-Flight Mass Spectrometry (GC-TOF MS) was used for Primary metabolites. For biogenic amines, separation and detections was achieved by Hydrophilic Interaction Chromatography (HILIC), followed by Quadrupole time-of-flight (QTOF) MS/MS. Lastly, for complex lipids, Charged Surface Hybrid (CSH™) C18 separation was followed with QTOF MS/MS for lipids. Extract preparation was also done at the same facility, from 1E+07 cells in each sample, from the same ones used for proteomic and RNA profiling (Figure 1). The cells were provided to the Metabolomics facility as frozen (at −80 °C) pellets. Detected species that could not be assigned to any compound were excluded from the analysis.

#### ANOVA-based computational approaches to identify differentially expressed biomolecules

For RNA samples, we used the TPM normalized values. For all other biomolecules, the input values we used were scaled-normalized for input values per sample. All the input and output datasets are shown in Table S1. To identify significant differences in the levels of biomolecules between any two points in the cell cycle we used the robust bootstrap ANOVA, via the *t1waybt* function, and the posthoc tests via the *mcppb20* function, of the *WRS2* R language package (Wilcox, 2011). The function is shown in File6, using as an example the ‘File2/sol_pa_anova spreadsheet. For this statistical analysis, the 24 cell size pools were grouped in 8 groups, for each of the approximately 5 fL increments in the cell size series (see Figure 1A).

#### SWATH-Mass spectrometry

The samples used to measure ribosomal protein abundances were from elutriated, diploid wild type BY4743 cells (see Key Resources Table). Once the cells reached the desired cell size, they were quenched with 100 µg/ml cycloheximide and 0.1% sodium azide. Cells were harvested from three independently elutriated cultures (5E+07 cells in each sample). The cells were re-suspended in a buffer containing 20 mM Tris·Cl (pH 7.4), 150 mM NaCl, 5 mM MgCl_2_, 1 mM DTT, 100 μg/ml cycloheximide, 1% ^v^/_v_ Triton X-100, and 25 U/ml Turbo DNase I, to a volume of 0.35 ml. Then, 0.2 ml of 0.5mm glass beads were added to each sample, and vortexed at maximum speed for 15 s, eight times, placing on ice for 15 s in between. The lysates were clarified by centrifuging at 5,000 rpm for 5 m, at 4 °C, and again for 5 m at 13,000 rpm at 4 °C. The supernatant was transferred to a 13×51 mm polycarbonate ultracentrifuge tube, underlaid with 0.90 ml of 1 M sucrose, and the ribosomes were pelleted by centrifugation in a TLA100.3 rotor (Beckman) at 100,000 rpm at 4 °C for 1 h. The protein pellets from three biological replicates for various time points during the cell cycle (40, 45, 50, 55, 60, 65, 70 and 75 fL) were then re-suspended in PBS, subjected to a Filter-Aided Sample Preparation (FASP) protocol tryptic digestion (Wisniewski *et al*., 2009), desalted using C-18 Ziptips, and analyzed by data-independent acquisition (DIA)/SWATH-mass spectrometry, as described previously (Schilling *et al*., 2017).

Briefly, samples were analyzed by reverse-phase HPLC-ESI-MS/MS using an Eksigent Ultra Plus nano-LC 2D HPLC system (Dublin, CA) with a cHiPLC system (Eksigent) which was directly connected to a quadrupole time-of-flight (QqTOF) TripleTOF 6600 mass spectrometer (SCIEX, Concord, CAN) (Christensen *et al*., 2018). After injection, peptide mixtures were loaded onto a C18 pre-column chip (200 µm × 0.4 mm ChromXP C18-CL chip, 3 µm, 120 Å, SCIEX) and washed at 2 µl/min for 10 min with the loading solvent (H_2_O/0.1% formic acid) for desalting. Subsequently, peptides were transferred to the 75 µm × 15 cm ChromXP C18-CL chip, 3 µm, 120 Å, (SCIEX), and eluted at a flow rate of 300 nL/min with a 3 h gradient using aqueous and acetonitrile solvent buffers.

For quantification, all peptide samples were analyzed by data-independent acquisition (Gillet *et al*., 2012), using 64 variable-width isolation windows (Collins *et al*., 2017; Schilling *et al*., 2017). The variable window width is adjusted according to the complexity of the typical MS1 ion current observed within a certain m/z range using a DIA ‘variable window method’ algorithm (more narrow windows were chosen in ‘busy’ m/z ranges, wide windows in m/z ranges with few eluting precursor ions). DIA acquisitions produce complex MS/MS spectra, which are a composite of all the analytes within each selected Q1 m/z window. The DIA cycle time of 3.2 s included a 250 ms precursor ion scan followed by 45 ms accumulation time for each of the 64 variable SWATH segments.

The DIA/SWATH data was processed with the Spectronaut™ software platform (Biognosys) for relative quantification comparing peptide peak areas among different time points during the cell cycle. For the DIA/SWATH MS2 data sets quantification was based on XICs of 6-10 MS/MS fragment ions, typically y- and b-ions, matching to specific peptides present in the spectral libraries. Significantly changed proteins were accepted at a 5% FDR (q-value < 0.05).

#### Immunoblot analysis

For protein surveillance, protein extracts were made as described previously (Amberg *et al*., 2006), and run on 4-12% Tris-Glycine SDS-PAGE gels. To detect TAP-tagged proteins with the PAP reagent, we used immunoblots from extracts of the indicated strains as we described previously (Blank *et al*., 2017). Loading was evaluated with an anti-Pgk1p antibody.

#### Comparison of the relative protein abundances in (Becher *et al*., 2018) and (Olsen *et al*., 2010)

For the datasets generated in human, HeLa cells, 0.5 h after nocodazole arrest, the data were from Table S1 in (Becher *et al*., 2018) and Supplementary Table_S1 in (Olsen *et al*., 2010). In the former study the authors reported the Log2-transformed ratios of the measured abundance over the median abundance of asynchronous cultures. For the (Olsen *et al*., 2010) proteins, the data were the isotopic ratios reported. In both cases, these values represented the corresponding protein abundances in that sample, among all the proteins identified in each sample in each study (see File7). To compare the rank order of the 3,298 proteins identified in common in the two studies, Spearman’s rank correlation rho (ρ) was estimated (ρ=0.09687857) with the spearman.test function of the *pspearman* R language package.

## SUPPLEMENTARY FIGURES

**FIGURE S1.**
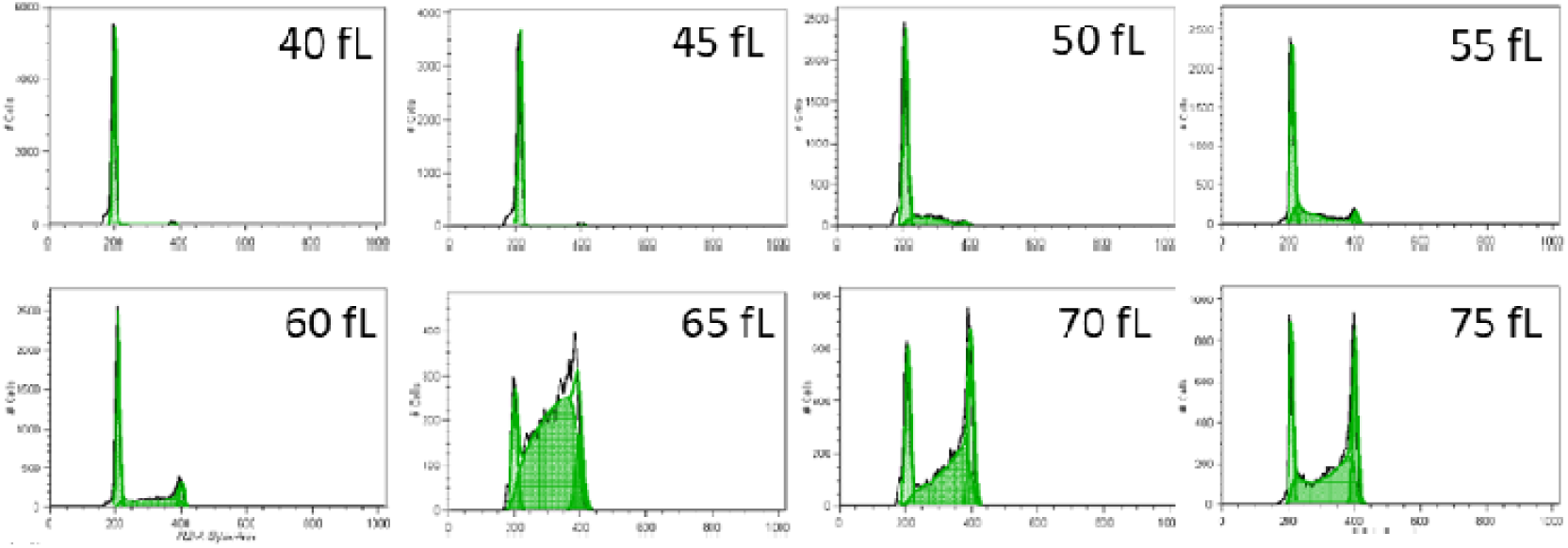
DNA content of samples spanning the cell size series from the elutriated samples. The DNA was measured with flow cytometry, as described in the Materials and Methods. On the y-axis of each histogram is number of cells and on the x-axis the fluorescence per cell.

**FIGURE S2.**
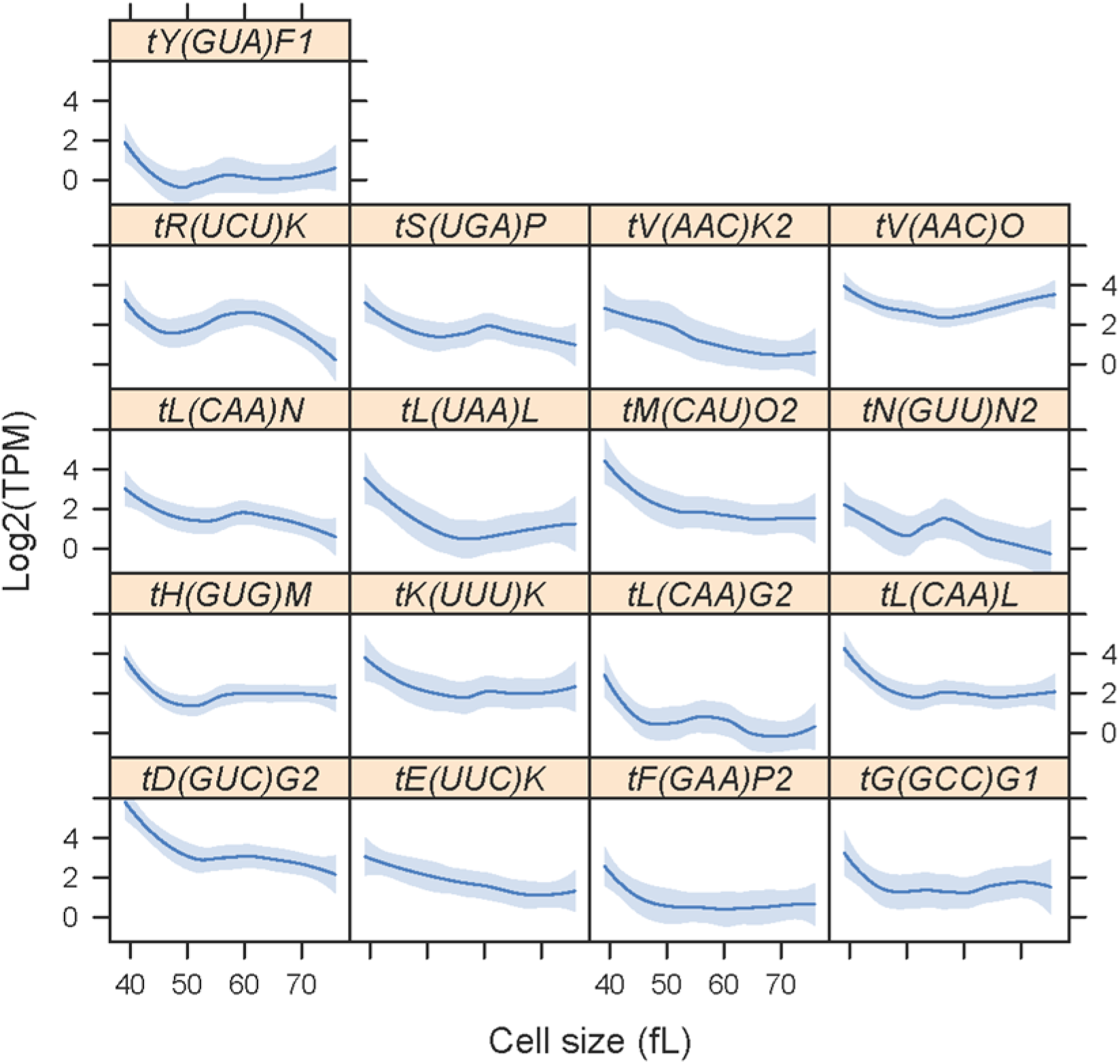
Levels of tRNAs, peaking early in the cell cycle. The tRNAs were from clusters 1 and 3 in Figure 2, with significantly different levels (p<0.05; Log2(FC)≥1) between any two points in the cell cycle, based on bootstrap ANOVA. Sequences corresponding to the tRNAs shown peaked in abundance at cell sizes from 40 to 50 fL. Cell size is shown on the x-axis (in fL), while the Log2-transformed ‘Transcripts Per Kilobase Million’ (TPM) values for each tRNA from all 24 data points are shown on the y-axis. Loess curves and confidence bands indicating the standard errors on the curve at a 0.95 level were drawn using the default settings of the panel.smoother function of the *latticeExtra* R language package.

**FIGURE S3.**
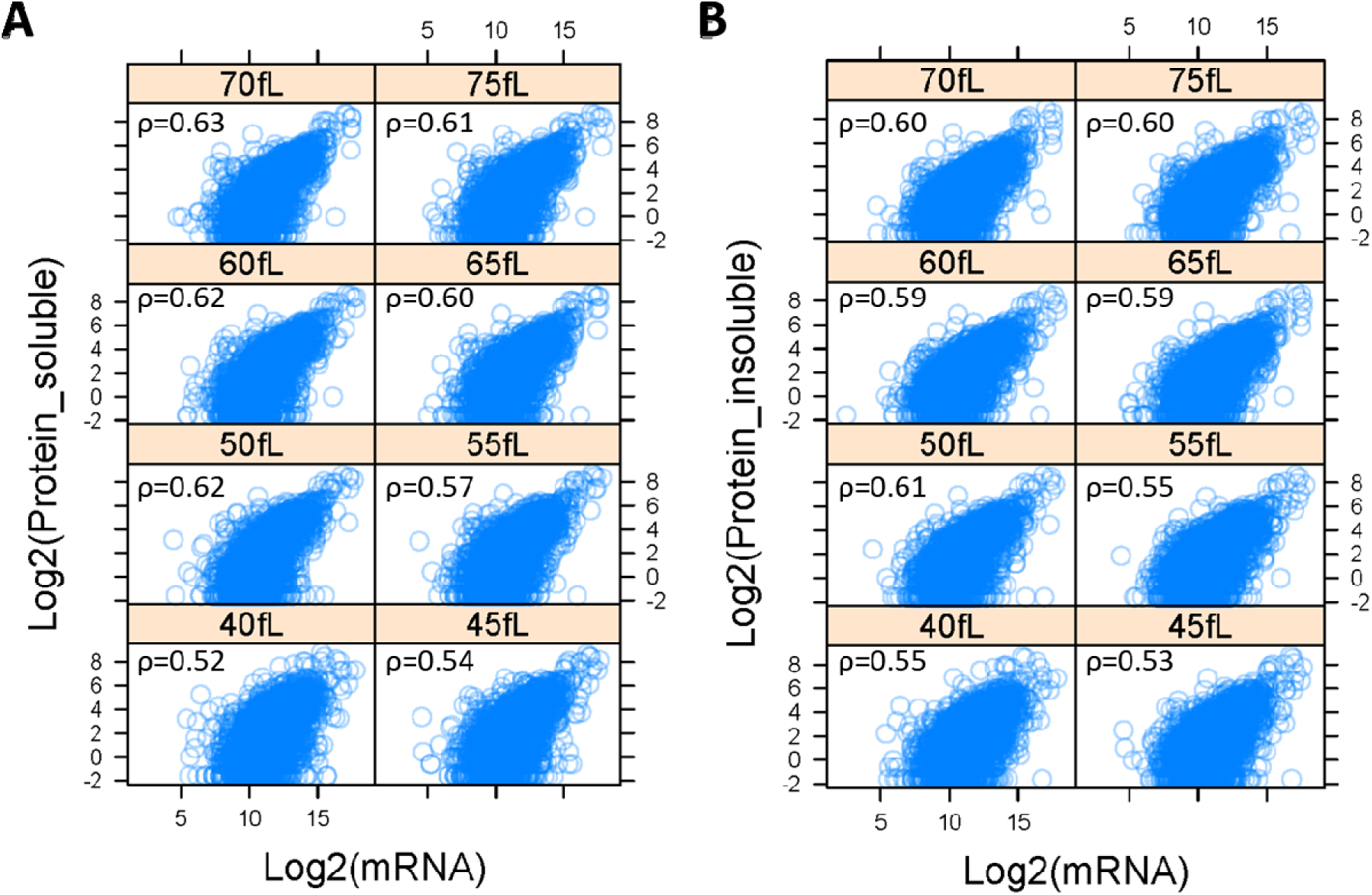
Transcriptome-proteome correlations. **A**, The spectral counts corresponding to the proteins identified in this study were averaged from the three biological replicates for each cell size pool we analyzed from the soluble fractions (from the ‘sol_psm’ dataset, see Table S1), and shown on the y-axis. On the x-axis are the RNA read counts from the corresponding loci (from the ‘rna_reads’ dataset, see Table 1). All values were Log2-transformed for display purposes. The Spearman correlation coefficients (ρ) shown in each case were calculated with the rcorr function of the *Hmisc* R language package. **B**, Similar analysis as in A, except that the input dataset for the spectral counts (y-axis) was from the insoluble proteome fractions (from the ‘pel_psm’ dataset, see Table S1).

**FIGURE S4.**
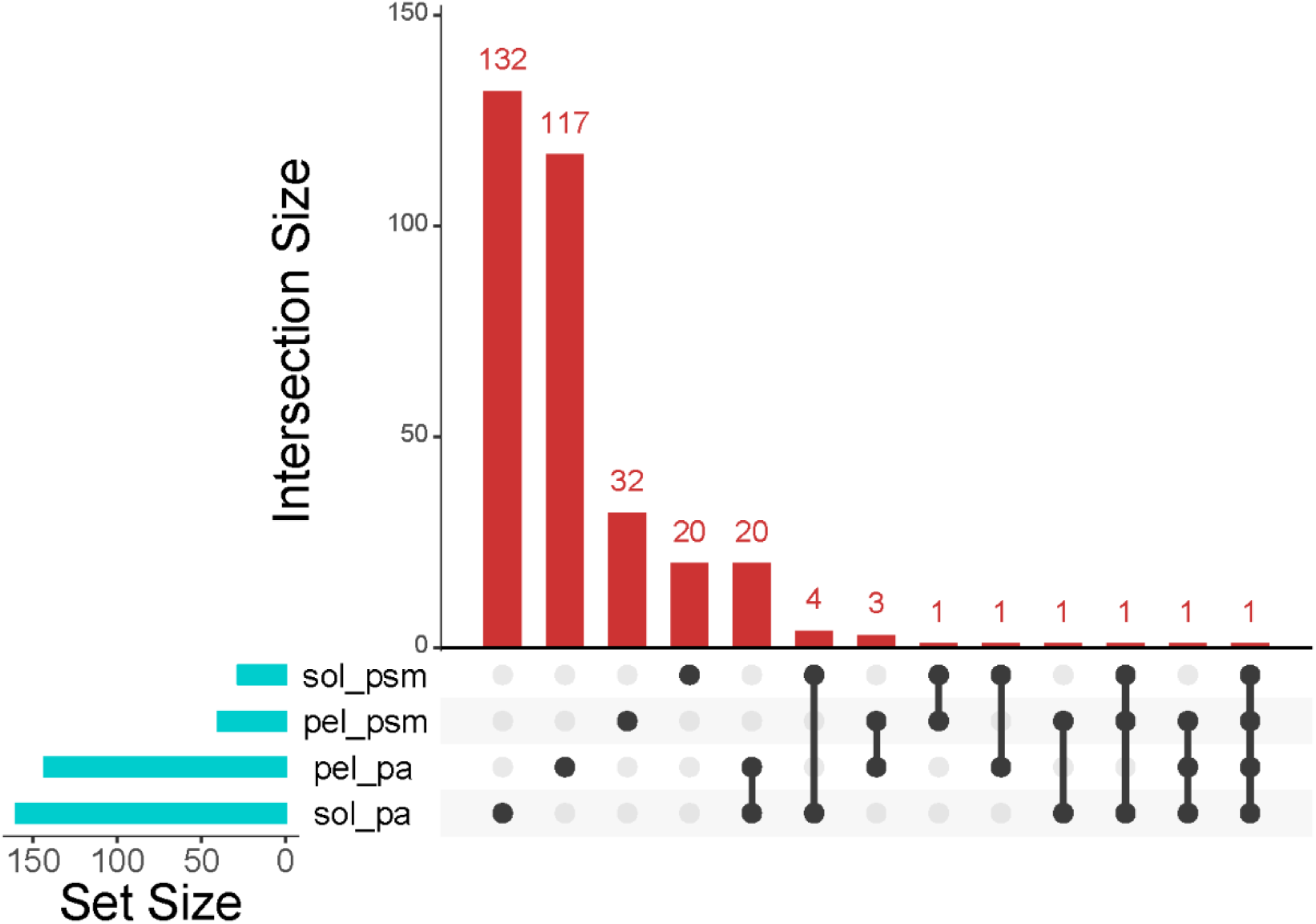
Overlap of protein datasets whose levels change in the cell cycle. Matrix layout for all intersections of the four ANOVA-identified sets containing proteins with significantly different levels (p<0.05; Log2(FC)≥1) between any two points in the cell cycle. The names of all proteins in each set are shown in File4/ Sheet: ‘proteins_sets’. The graph was drawn with the UpSet R language package.

**FIGURE S5.**
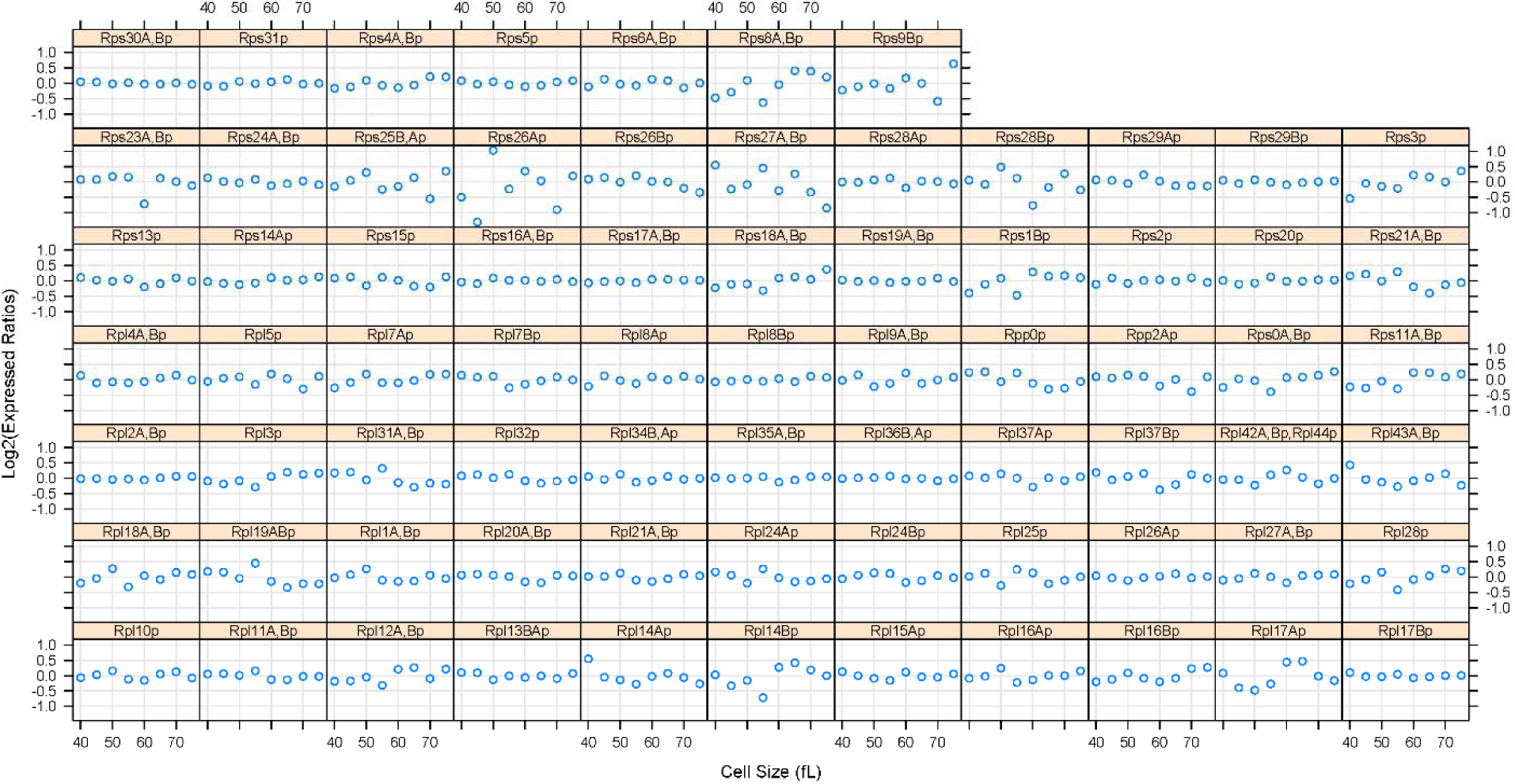
Ribosomal protein abundance in ribosomes is not periodic in the cell cycle. The levels of each ribosomal protein (see Figure 4) detected were normalized against the sum of all ribosomal proteins detected in that sample, and displayed as Log2-transformed expressed ratios (y-axis), while cell size (in fL) is on the x-axis. In none of the few cases (e.g., Rps8,9,26,27,28p; Rpl14,17Ap) where the abundance of the ribosomal protein in question appeared to fluctuate somewhat in the cell cycle the changes were periodic (FDR>0.05), and these changes likely reflect experimental error in the quantification.

**FIGURE S6.**
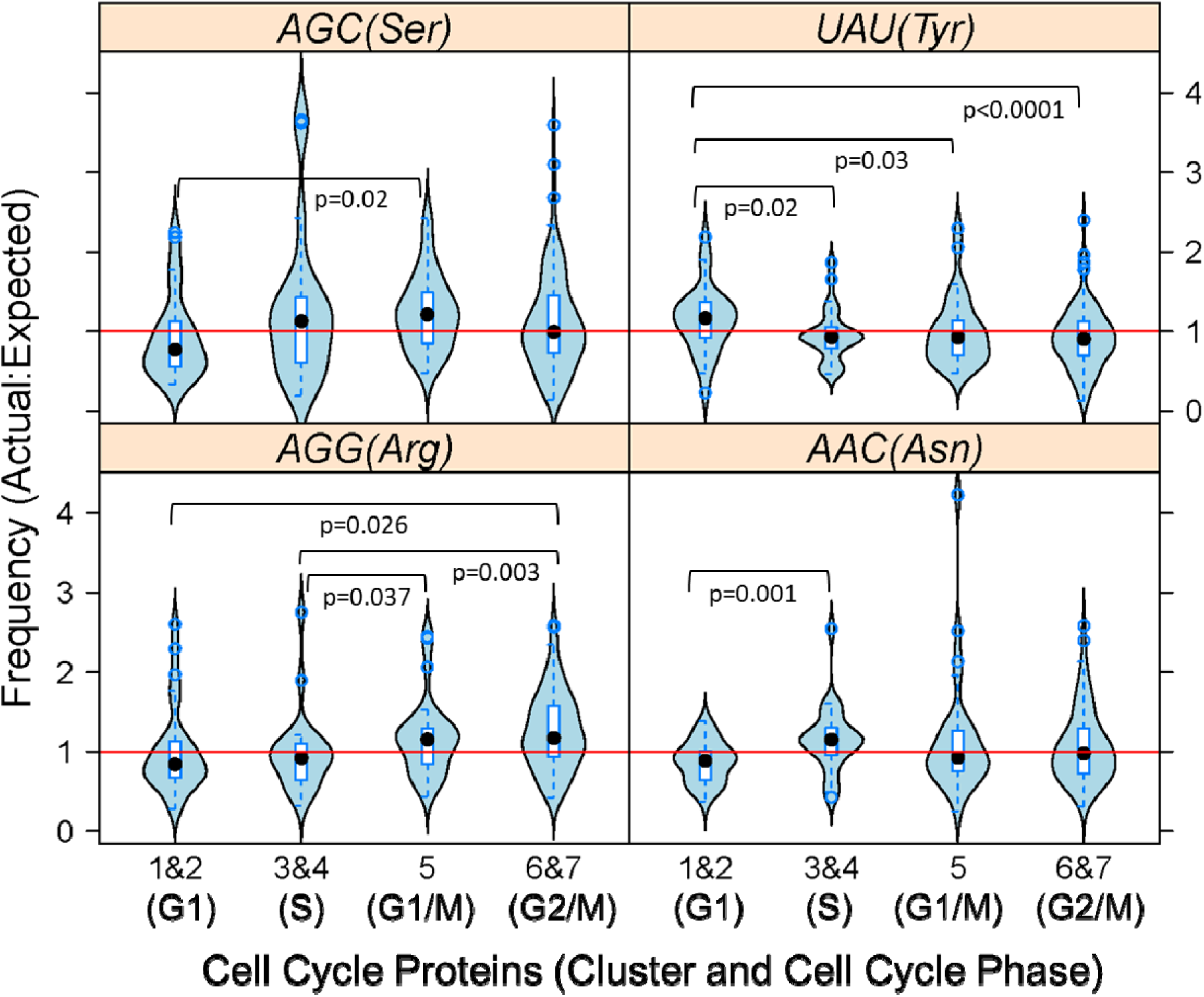
Little, if any, evidence for cell cycle-dependent changes in codon usage. From the 333 cell cycle-regulated proteins shown in Figure 3, we selected the ones who were not identified as ubiquitylated by (Swaney *et al*., 2013), and whose corresponding mRNA levels were not changing (from Figure 2). These proteins were further grouped according to their cell cycle expression pattern (peaking in G1: in clusters 1&2 (n=29); peaking in S: in clusters 3&4 (n=24): peaking in G1/M: in cluster 5 (n=29); peaking in G2/M: in clusters 6&7 (n=90)). For each codon in each mRNA encoding each of these proteins, we obtained the ratio of the actual to expected usage, based on (Tumu *et al*., 2012). These values are displayed as violin plots, for the four codons shown that there were statistically significant differences between the groups for each codon (based on bootstrapped ANOVA: p<0.05). For differences between groups in each codon, the p-values shown were obtained from posthoc statistical tests, using the mcppb20 function of the *WRS2* R language package. The red horizontal lines indicate equal actual:expected codon usage in each case.

